# The function of α-Catenin mechanosensing in tissue morphogenesis

**DOI:** 10.1101/2021.09.08.459511

**Authors:** Luka Sheppard, Ulrich Tepass

**Author notes:** **Correspondence should be addressed to:** Ulrich Tepass, Department of Cell and Systems Biology University of Toronto, 25 Harbord Street, Toronto, Ontario M5S 2M6 Canada, **Phone** (all authors), **Emails**.

## Abstract

α-catenin couples the cadherin-catenin complex to the actin cytoskeleton. The mechanosensitive α-catenin M region undergoes conformational changes upon application of force to recruit binding partners. Here, we took advantage to the tension landscape in the Drosophila embryo to define three different states of α-catenin mechanosensing in support of cell adhesion. Low, medium, and high tension contacts showed α-catenin M region-dependent low, medium, and high levels of Vinculin and Ajuba recruitment. In contrast, Afadin/Canoe acts in parallel to α-catenin at bicellular low and medium tension junctions, but requires an interaction with α-catenin for its tension-sensitive enrichment at high-tension tricellular junctions. Individual M region domains make complex contributions to cell adhesion through their impact on binding partner recruitment, and redundancies with the function of Afadin/Canoe. Our data argue that α-catenin and its interaction partners are part of a cooperative and partially redundant, mechanoresponsive network that supports AJs remodelling during morphogenesis.

## Introduction

Adherens junctions (AJs) resist actomyosin generated forces to change cell and tissue shape while maintaining tissue cohesion. Dynamic AJs are disassembled and rebuilt in concert with cell contact changes during tissue morphogenesis. Combining both dynamic and stable aspects of AJs requires the interplay of a complex molecular machinery. Significant progress has been made in our understanding of the molecular architecture of AJs since the original discovery of cadherin adhesion molecules (Harris and Tepass, 2010; Maître and Heisenberg, 2013; Takeichi, 2014; 2018; Pinheiro and Bellaïche, 2018). However, much remains to be learned about how AJs operate during specific morphogenetic movements and in particular how AJs associate with the actin cytoskeleton to support cell shape changes and cell movements in tissues such as epithelia.

As tissues adopt new forms during morphogenesis, cell contacts are subjected to contractile forces to elicit coordinated cell shape changes and cell rearrangements. The ability of AJ components to respond to force is believed to be key to maintaining cohesion and tissue integrity during the cell-cell contact changes that underpin morphogenesis (Lecuit and Yap, 2015; Ladoux et al., 2015; Pinheiro and Bellaïche, 2018; Charras and Yap, 2018; Clarke and Martin, 2021). The cadherin-catenin complex (CCC) physically couples the cytoskeleton of neighboring cells and in response to force — generated often by actomyosin contraction — strengthens adhesive interactions (Le Duc et al., 2010; Yonemura et al., 2010). Extracellular cadherin domains experience force-induced conformation changes, increasing the strength of trans-interactions between cadherins of neighboring cells (Leckband and De Rooij, 2014; Pinheiro and Bellaïche, 2018). The transmission of force through cadherins and the integrity of AJs requires the intracellular physical link to the actin cytoskeleton provided by α-catenin (Rimm et al., 1995; Desai et al., 2013; Buckley et al., 2014). α-catenin is thought to be the central mechanosensor of the CCC (Yonemura et al., 2010; Ishiyama and Ikura, 2012; Leckband and De Rooij, 2014; Ladoux et al., 2015; Angulo-Urarte et al., 2020). Under the application of force, conformational changes within α-catenin lead to enhanced actin binding, the downstream regulation of actin networks, and recruitment of additional AJ components (Yonemura et al., 2010; Choi et al., 2012; Rangarajan and Izard, 2012; Twiss et al., 2012; Huveneers et al., 2012; Barry et al., 2014; Mège and Ishiyama, 2017; Ishiyama et al., 2018; Sarpal et al., 2019; Alégot et al., 2019). Mutational analysis of α-catenin has demonstrated its essential role for epithelial cell adhesion similar to E-cadherin (Ecad) and β-catenin (Armadillo [Arm] in Drosophila) in a number of animal species and diverse tissues (Torres et al., 1997; Kofron et al., 1997; Costa et al., 1998; Schepis et al., 2012; Sarpal et al., 2019; Clarke et al., 2019). However, the function of α-catenin mechanosensing in tissue morphogenesis remains largely unexplored.

The two mechanosensory domains of α-catenin are the central M region and the C-terminal actin-binding domain (ABD) (Fig. 1A,B). α-catenin directly binds F-actin through the ABD as a catch-bond, whereby under increasing tension, the strength of this bond, the bond lifetime, increases up to a threshold (Buckley et al., 2014). Force-induced conformational change in the ABD enhances direct F-actin binding (Ishiyama et al., 2018; Xu et al., 2020). In contrast, force-induced conformational changes in the central M region of α-catenin cause the recruitment of other F-actin binding proteins, such as Vinculin and Ajuba, which is thought to reinforce adhesion (Mège and Ishiyama, 2017; Angulo-Urarte et al., 2020).

**Figure 1.**
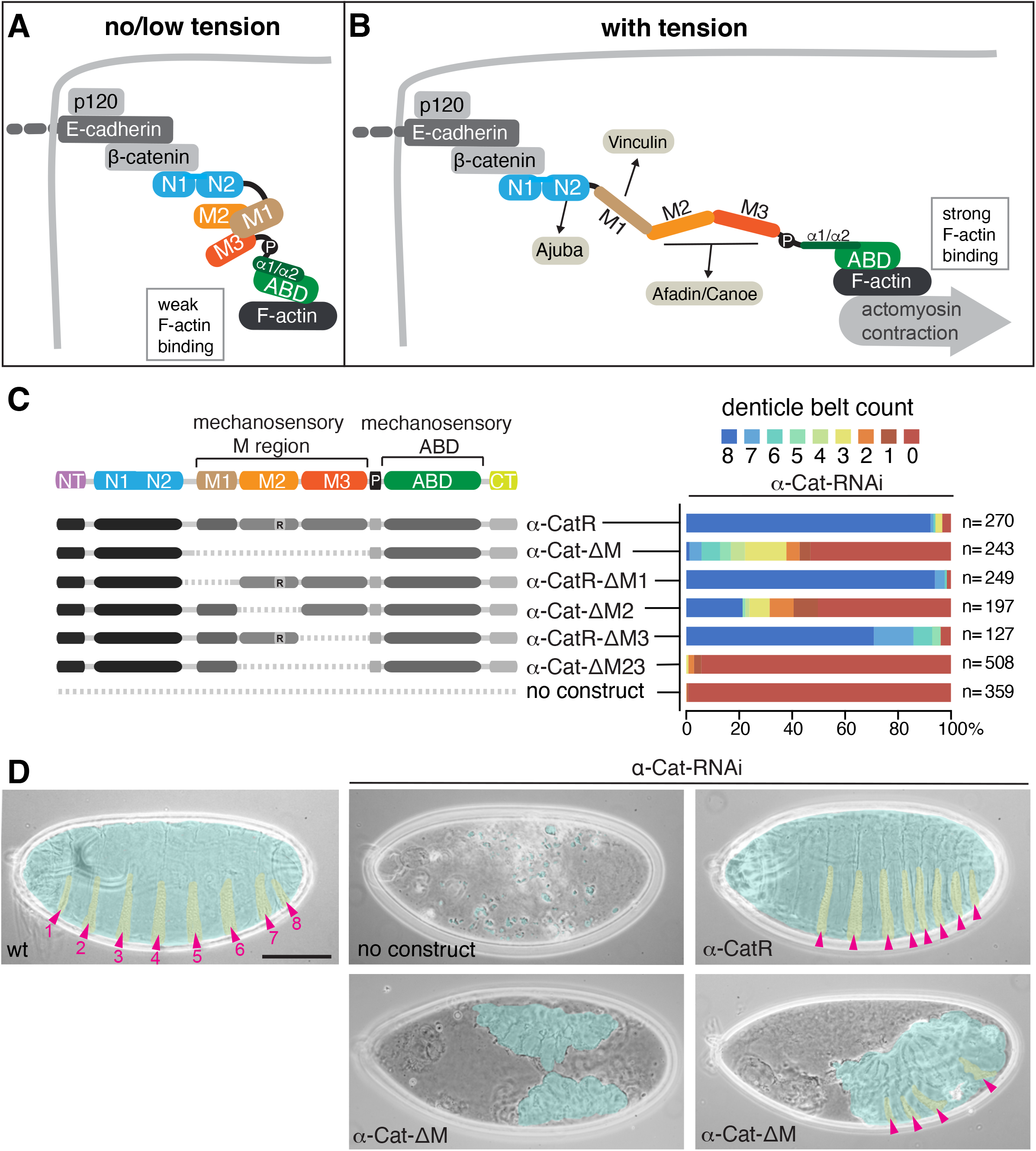
The α-Cat M region is required for epithelial integrity during embryogenesis. **(A,B)** Schematic of α-Cat under no/low and high tension. See text for further discussion. **(C)** Quantification of cuticle defects of embryos expressing α-CatR or α-Cat deletion constructs in an α*-Cat-RNAi* background. **(D)** Representative cuticle images of embryos of the indicated genotypes. False color shading of cuticle in blue and denticle belts in yellow. Pink arrowheads point to abdominal denticle belts that were used for quantification in (C). Scale bar, 100μm.

Here we assessed the function of the M region and its three domains (M1, M2, and M3) in embryonic morphogenesis. M1, M2 and M3 consist of α-helical bundles that without tension form a closed conformation supported by a dynamic electrostatic network produced by salt-bridges between these bundles (Ishiyama et al., 2013; Li et al., 2015). Application of tension breaks the salt bridges, exposing cryptic binding sites (Fig. 1A,B). The M1 domain unfurls and binds Vinculin (Vinc), recruiting Vinc to the cell membrane (Yonemura et al., 2010; Ishiyama et al., 2013; Barry et al., 2014; Yao et al., 2014; Kim et al., 2015; Maki et al., 2016; Seddiki et al., 2018). Moreover, the junctional recruitment of Ajuba (Jub) is regulated by mechanosensing through the M region, with M1 preventing Jub recruitment to α-catenin under low tension (Alégot et al., 2019; Sarpal et al., 2019). A second conformational change in the M region was reported for the M2 and M3 domains that may involve an increase in the angle between M2 and M3 or the partial or complete unfurling of these α-helical bundles, which could impact the interaction between α-catenin and Afadin (Pokutta et al., 2002; Ishiyama et al., 2013; Yao et al., 2014; Li et al., 2015; Matsuzawa et al., 2018; Sakakibara et al., 2020).

We examined the role of the M region in supporting adhesion during Drosophila embryonic morphogenesis, and how the M region cooperates with the α-Catenin (α-Cat) interactors Vinc, Jub, and Canoe (Cno, the Drosophila ortholog of Afadin) in maintaining tissue integrity as cells undergo shape changes and rearrangements. Contrary to expectations, our recent work has shown that an α-Cat constructs which lack the entire M region can fully replace endogenous α-Cat in the development of the wing disc epithelium (Sarpal et al., 2019). Deletion of individual M region domains could also support normal epithelial integrity of the disc epithelium, and the ovarian follicular epithelium (Desai et al., 2013; Sarpal et al., 2019). However, deletion of M1 interfered with the normal regulation of tissue growth. Tension sensing through M1 is required to prevent the constitutive recruitment of Jub to α-Cat. Junctional Jub forms a complex with the Hippo pathway kinase Warts to downregulate the Hippo pathway and promote growth through activation of Yorkie (Rauskolb et al., 2014; Alégot et al., 2019; Sarpal et al., 2019). This mechanism allows tissue tension to regulate growth and is the only example to date demonstrating a function for α-catenin M region mechanosensing in a whole organism. In particular, how M region mechanosensing contributes to cell adhesion in tissue morphogenesis remains an open question. The situation is further complicated by the lack of evidence for Vinc as an important player in cell adhesion *in vivo*. Several tissue culture studies have suggested that the tension-dependent binding of Vinc to the unfurled M1 domain makes an important contribution to adhesion mediated by the CCC (Yonemura et al., 2010; Le Duc et al., 2010; Huveneers et al., 2012; Twiss et al., 2012; Seddiki et al., 2018). In striking contrast, analysis of *Vinc* mutant animals in *C. elegans* (Barstead and Waterston, 1991), Drosophila (Alatortsev et al., 1997; Maartens et al., 2016), zebrafish (Han et al., 2017), and mouse (Xu et al., 1998) does not support a significant role of Vinc in cadherin-based cell adhesion. Similarly, Jub null mutants in the mouse, zebrafish and Drosophila complete embryogenesis with only subtle defects (Pratt et al., 2005; Witzel et al., 2012; Razzell et al., 2018) consistent with a minor role in cell adhesion, at best.

The only α-catenin M region binding partner with substantive adhesion defects, although weaker than those resulting from the loss of α-catenin, is Afadin/Cno. M3 contains a minimal Afadin binding site, with the exposed M2 domain also contributing to this interaction (Pokutta et al., 2002; Sakakibara et al., 2020). In Drosophila, *cno* maternal zygotic mutants cause defects in mesoderm invagination (Sawyer et al., 2009) and tears in the ectodermal epithelium as it undergoes convergent extension (Sawyer et al., 2009; Yu and Zallen, 2020). *cno* zygotic mutants have dorsal closure defects (Jürgens et al., 1984; Takahashi et al., 1998; Boettner et al., 2003; Choi et al., 2011) similar to some *α-Cat* zygotic mutants (Sarpal et al., 2012; Jurado et al., 2016). In mouse *afadin* mutants, strong defects arise after implantation where the neural fold becomes a mass of disorganized cells, embryos remain flat and short and lethality ensues (Ikeda et al., 1999; Zhadanov et al., 1999). Actomyosin cables in cells of the extending germband of Drosophila *cno* mutants are more diffuse and detached from the membrane, making unproductive contractions (Sawyer et al., 2011). In cell culture, a mutant α-catenin isoform with an open conformation of the M region showed increased Afadin recruitment (Matsuzawa et al., 2018), and the α-catenin binding site on Afadin is required for its mechanosensitive enrichment (Sakakibara et al., 2020). We wondered therefore whether Cno is acting as the effector of the M region to support adhesion in Drosophila. The role of α-catenin M region domains in Afadin/Cno recruitment and function in adhesion has not been explored in an animal model.

The comparatively mild to moderate defects produced by the loss of Vinc, Jub, and Afadin/Cno led us to question (i) whether, and if so, how the α-Cat M region contributes to adhesion and embryonic morphogenesis via recruitment of these binding partners in response to force, and (ii) whether there is redundancy between α-Cat binding partners which would explain the moderate to subtle phenotypes that result from the loss of Vinc, Jub, and Cno compared to the striking loss of epithelial integrity that is caused by the loss of α-Cat. We found that the M region is required for cell adhesion during early Drosophila embryogenesis, in particular the M2 domain, at contacts that experience higher tension. Our data suggest three distinct tension states read by α-Cat mechanosensing that cause the differential recruitment of Vinc, Jub, and Cno to enhance adhesion. Surprisingly, ectopic exposure of the M1 domain was more deleterious than removal the M region entirely. This effect is not due to the recruitment of Vinc, but likely to the M1-dependent regulation of Jub recruitment to AJs. Our findings also support the conclusion that Cno promotes cell adhesion in a parallel pathway to the CCC except at tricellular junctions (TCJs) where the mechanosensitive enrichment of Cno (Yu and Zallen, 2020) depends on the M2 and M3 domains of α-Cat. Our work provides evidence of a robust network of cooperative and redundant mechanosensitive interactions at AJs that support tissue morphogenesis.

## Results

### The M region of α-Cat is required for epithelial integrity during embryonic morphogenesis

We have previously analyzed the ability of mutant α-Cat proteins to substitute for endogenous α-Cat in several different Drosophila tissues including the embryonic head epidermis, the follicular epithelium of the ovary, and the epithelium of the wing imaginal disc (Sarpal et al., 2012; Desai et al., 2013; Escobar et al., 2015; Ishiyama et al., 2018; Sarpal et al., 2019). Expression of an α-Cat construct that lacks the entire M region (α-Cat-ΔM) could fully support epithelial development of the wing discs, which developed into a normal adult wing (Sarpal et al., 2019). Further analysis of α-Cat-ΔM showed that it provided a strong rescue of *α-Cat* mutant cells of the follicular epithelium (Fig. S1), supported development of the head epidermis and prevented embryonic lethality of most zygotic *α-Cat* null mutant animals (Sarpal et al., 2019). In contrast to the widely considered model that M region-based mechanosensing enhances adhesion in response to mechanical force (Leckband and De Rooij, 2014; Mège and Ishiyama, 2017; Angulo-Urarte et al., 2020), our *in vivo* data at this point assigned only a minor or no essential role to the M region in cell adhesion and epithelial development. We therefore wondered whether M region function is most relevant during developmental periods of vigorous morphogenesis when cell contacts experience high levels of force. Drosophila gastrulation represents a period of development that condenses several large-scale morphogenetic movements into a short ∼2 hour time window. These movements entail the invagination of the mesoderm and endoderm (Martin, 2020), cell intercalations that drive germband extension (Paré and Zallen, 2020), the ingression of neural stem cells (Simões et al., 2017; An et al., 2017), and three rounds of cell division with a ∼40 minute cell cycle time (Foe, 1989).

To assess M region function in early embryos, we utilized a setup in which the crucial maternal contribution to *α-Cat* expression (Sarpal et al., 2012) is removed through RNA interference (Fig. S2A,B) and constructs are expressed that are resistant to the shRNA used (Sarpal et al., 2019). Our constructs are recruited to AJs (Fig. S2C) and are expressed at close to normal levels (Desai et al., 2013; Sarpal et al., 2019). Resistant constructs contain a mutated shRNA target site that preserves the amino acid sequence located in the M2 domain (denoted by ‘R’ ; e.g. α-CatR or α-CatR-ΔM1, with mutant α-Cat isoforms collectively referred to as α-CatX) or carry a deletion of M2 (Fig. 1C). Maternally driven *α-Cat-RNAi* caused a dramatic phenotype similar to that previously reported for the loss of Arm (Cox et al., 1996) or Ecad (Tepass et al., 1996), leading to a terminal phenotype with a highly fragmented epidermal/cuticle layer (Fig. 1D). We quantified the cuticle defects, which are indicative of adhesion defects, by counting the number of intact abdominal denticle belts. *α-Cat-RNAi* embryos had 0 intact denticle belts. *α-Cat-RNAi* embryos expressing α-CatR had all 8 denticle belts restored in >90% of animals. In contrast, *α-Cat-RNAi α-Cat-ΔM* animals showed partial restoration of epidermal integrity compared to *α-Cat-RNAi,* but virtually all animals had denticle belt defects with more than half of embryos producing no denticle belts (Fig. 1C,D). These results demonstrate a substantive requirement of the M region in maintaining epithelial integrity in the Drosophila embryo.

To investigate the function of the M region in early morphogenesis, live embryos were observed during gastrulation with a focus on mesoderm invagination and ectodermal integrity during germband extension. Mesoderm invagination failed in *α-Cat-RNAi* embryos (Fig. 2A, Fig. S2B). As previously described (Martin et al., 2010), at the onset of mesoderm invagination actomyosin contraction overpowers residual adhesion in *α-Cat-RNAi* embryos. Cells round up and come apart from each other, with plasma membrane tethers formed between few remaining AJ puncta (Fig. S2D). *α-Cat-RNAi α-CatR* embryos showed normal mesoderm invagination. In contrast, *α-Cat-RNAi α-CatR-ΔM* embryos displayed a range of defects that were classified according to the degree of successful invagination. ∼40% of embryos showed normal mesoderm invagination where the ventral midline sealed along the entire anterior-posterior axis, whereas the remaining ∼60% of embryos showed either a partial closure or a complete failure to close the ventral midline (Fig. 2A).

**Figure 2.**
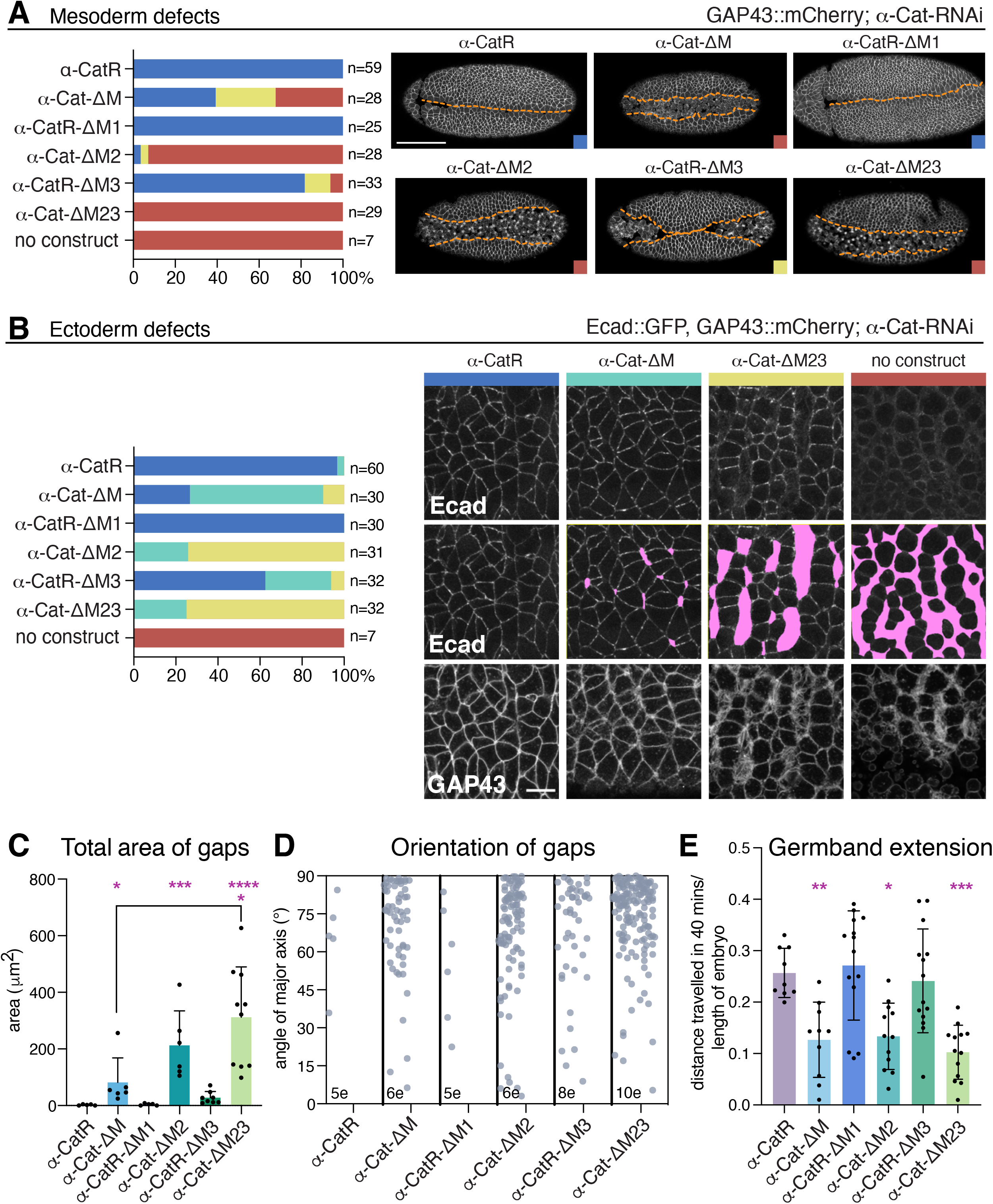
The α-Cat M region is essential for adhesion during mesoderm invagination and germband extension. **(A)** Quantification of mesoderm invagination defects in embryos of indicated genotype. Color coded example images are given at right. Orange lines indicate edges of mesoderm. Red - open ventral furrow; Yellow - partially fused ventral furrow; Indigo - >90% midline fusion. Scale bar; 100μm. **(B)** Quantification of defects in the lateral ectoderm during germband extension in embryos of indicated genotype. Classification used: Indigo - wild type, very few gaps seen; Cyan - small gaps; Yellow - large gaps or tears; Red – few if any identifiable AJs. Pink areas represent regions of gaps where apical junctions have lost contact. Scale bar, 10μm. **(C)** Area of gaps in *α-Cat-RNAi α-CatX* embryos. **(D)** Plot showing the angle of the major axis of ellipses fitted to each gap in *α-Cat-RNAi α-CatX* embryos. N=number of embryos(e) **(E)** Extension of the germband observed over 40 minutes in *α-Cat-RNAi α-CatX* embryos. For (C) and (E) significance given by ordinary one-way ANOVA, (**** = P <0.0001, ***= P <0.0002, **=P<0.0021, *=P <0.0332).

As the germband starts to extend, ectodermal cells in *α-Cat-RNAi* embryos lose adhesion and large gaps appear between the apical domain of cells (Fig. 2B). Depletion of the maternal contribution of α-Cat eventually leads to complete cell dissociation and epithelial collapse (Cavey et al., 2008; Martin et al., 2010; Rauzi et al., 2010; Fernandez-Gonzalez and Zallen, 2011; Wang et al., 2013; Levayer and Lecuit, 2013a; Eritano et al., 2020). Similar to the mesoderm, plasma membrane tethers are seen connecting cells across gaps (Fig. S2E) (Fernandez-Gonzalez and Zallen, 2011). *α-Cat-RNAi α-CatR* embryos displayed normal cell contacts in the ectoderm, whereas *α-Cat-RNAi α-CatR-ΔM* embryos showed gaps in the epithelium (Fig. 2B,C). Neighboring apical domains became separated, leading to epithelial gaps or tears that extend predominantly along the dorsal-ventral (DV) axis (Fig. 2D). Membrane tethers can be observed spanning these gaps (Fig. 2B). These defects were associated with a marked reduction in germband extension compared to *α-Cat-RNAi α-CatR* controls (Fig. 2E). Thus, the M region makes an essential contribution to maintaining adhesion during mesoderm invagination and germband extension and its loss compromises these processes.

### Exposure of the M1 domain is more deleterious to development than removal of the whole M region

We next assessed the performance of α-Cat constructs that lacked individual M region domains (α-CatR-ΔM1, α-Cat-ΔM2, α-CatR-ΔM3) or both M2 and M3 (α-Cat-ΔM23). α-CatR-ΔM1 behaved like full-length α-CatR when we examined the cuticle (Figs. 1C and S2F), mesoderm invagination (Fig. 2A), the ectodermal epithelium (Fig. 2B-D), and germband extension (Fig. 2E), suggesting that the M1 domain makes no essential contribution to cell adhesion in these tissues. Surprisingly, expression of α-Cat-ΔM23 was much less capable of ameliorating the *α-Cat-RNAi* phenotype than α-Cat-ΔM. *α-Cat-RNAi α-Cat-ΔM23* embryos showed little improvement of the cuticle defects seen with *α-Cat-RNAi* (Figs. 1C and S2F) and failed to rescue mesoderm invagination (Fig. 2A). Most embryos displayed prominent de-adhesion defects in the ectoderm, though these are not as severe as with *α-Cat-RNAi* alone (Fig. 2B-D), and germband extension was substantially reduced (Fig 2E). Rescue by α-Cat-ΔM2 showed minor improvements when compared to α-Cat-ΔM23, whereas expression of α-CatR-ΔM3 showed a much better rescue than α-Cat-ΔM2 (Figs 1C, 2, and S2F). This suggests that M2 is the most important domain within the M region for supporting cell adhesion, with M3 making a minor contribution. Notably, the poor rescue activity seen with α-Cat-ΔM23 compared to α-CatR-ΔM suggests that the unfurled M1 domain, which is retained in α-Cat-ΔM23, negatively regulates cell adhesion.

### The M region contributes to cell adhesion at medium and high tension cell contact sites

The ectoderm during germband extension displays an asymmetric distribution of non-muscle myosin II, characterized by an enrichment of myosin at vertical cell edges (oriented along the DV axis) and even more so at tricellular junctions (TCJs) (Bertet et al., 2004; Zallen and Wieschaus, 2004; Fernandez-Gonzalez et al., 2009; Tetley et al., 2016; Vanderleest et al., 2018). For the purpose of our discussion and reflecting the distribution of myosin, we distinguish between low-tension horizontal bicellular junctions (BCJs), medium-tension vertical BCJs, and high-tension TCJs (vertices). Contractions of myosin at vertical edges either in multicellular rosettes or at bicellular contacts are important drivers of cell intercalation required for germband extension (Zallen and Wieschaus, 2004; Blankenship et al., 2006; Zallen and Blankenship, 2008; Fernandez-Gonzalez et al., 2009; Tetley et al., 2016; Vanderleest et al., 2018) and neuroblast ingression (Simões et al., 2017). We noticed that gaps or tears in the epithelium of embryos expressing M region deletions were commonly found at the centers of rosettes or at vertices, and were elongated along the DV axis (Fig. 2B,D). These findings further suggest that the M region and its individual domains, except for M1, strengthen adhesion and that medium and high tension cell contact sites are more susceptible to loss of adhesion than other cell contacts.

To further characterize these adhesion defects, we analyzed the actomyosin cytoskeleton using the endogenously YFP-tagged myosin heavy chain. The expression of any M region deletion in an *α-Cat-RNAi* background led to increased total myosin signal (Fig. 3A and E), while the enrichment of myosin to vertical edges was consistently observed (Fig. 3A). As myosin is recruited by tension (Fernandez-Gonzalez et al., 2009) this suggests that the asymmetric distribution of tension in the ectoderm is not strongly affected in embryos expressing M region deletions. Although membrane association of myosin was preserved even with a poor rescue of adhesion as in *α-Cat-RNAi α-Cat-ΔM23* embryos, separation of apical domains along the DV axis led to the apparent splitting of supracellular myosin cables (Fig. 3B). In embryos with stronger defects, as gaps open between ectodermal cells, myosin accumulates in large puncta at the free apical cell edges (Fig. 3B).

**Figure 3.**
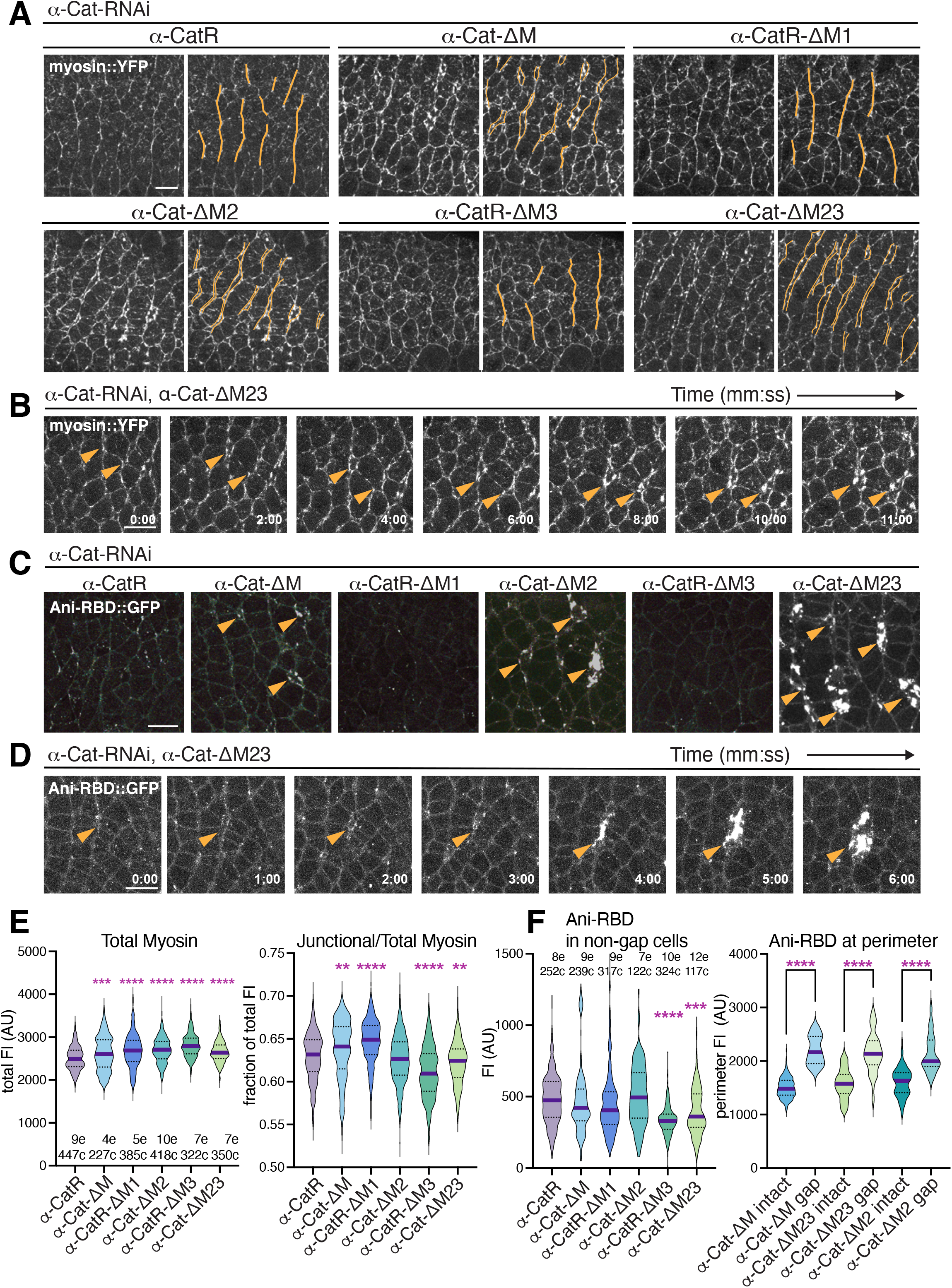
Myosin distribution and Rho1 activity in embryos expressing α-Cat M region deletions. **(A)** Representative images of ectoderm during germband extension of *α-Cat-RNAi α-CatX* embryos. Myosin cables are highlighted with orange lines in the right duplicate images. **(B)** Series of stills from a live *α-Cat-RNAi α-Cat-ΔM23* embryo showing gaps forming at vertical edges (arrowheads) with accumulations of myosin at gap perimeters. **(C)** Stills from live *α-Cat-RNAi α-CatX* embryos expressing the Rho1 activity probe, Ani-RBD::GFP. Note accumulation of Ani-RBD::GFP in gap areas (arrowheads). **(D)** Series of stills from a live *α-Cat-RNAi α-Cat-ΔM23* embryo showing enrichment of Ani-RBD::GFP at a forming gap between cells (arrowheads). Cell contacts first split and subsequently Ani-RBD signal increases. **(E)** Quantification of myosin *α-Cat-RNAi α-CatX* embryos. **(F)** Quantification of Ani-RBD::GFP signal at cortices of cells which are not in contact with a gap. Average FI of Ani-RBD::GFP signal at the perimeter of cells is compared to the perimeter of gaps. For (E) and (F) significance is calculated by ordinary one-way ANOVA (**** = P <0.0001, ***= P <0.0002, **=P<0.0021). N= number of cells (c), and embryos (e).. Scale bar in (A-D), 10μm.

Gaps in the ectoderm of *α-Cat-RNAi α-CatX* embryos could also be visualized by an enhanced signal of the active Rho1 probe Ani-RBD. Activation of Rho1 causes the phosphorylation of myosin regulatory light chain by Rho kinase, and hence myosin contraction, with enrichments of activated Rho1 seen at sites of high actomyosin contractility in the germband (Munjal et al., 2015; Martino et al., 2018). Loss of apical cell contact was accompanied by the formation of large puncta of Ani-RBD (Fig. 3C,D,F). These observations are reminiscent of the accumulation of active Rho1 and myosin at wound margins (Abreu-Blanco et al., 2014; Rothenberg and Fernandez-Gonzalez, 2019), and Rho flares which repair breaches in the epithelial barrier during Xenopus development (Stephenson et al., 2019). The appearance of membrane tethers (Fig. 2B) and the accumulation of active Rho1 and myosin (Fig. 3) confirmed the loss of cell-cell contacts within the germband, particularly at medium and high tension cell edges or vertices, for embryos expressing deletions that remove the M2 domain (Figs. 2B,C and 3C). Moreover, α-CatR-*Δ*M3 and α-Cat-*Δ*M23 expressing embryos showed comparatively lower levels than α-CatR in both cortical Ani-RBD signal and the proportion of junctional myosin (Fig. 3E,F). In contrast, a higher fraction of junctional myosin was seen when M1 was deleted (Fig. 3E). Together, these findings suggest that the M region contributes to the regulation of junctional actomyosin.

### M2/3 and M1 play opposing roles in regulating AJs

To further characterize the impact of the M region on cell adhesion we examined the junctional distributions of Ecad and the α-Cat binding partners Vinc and Jub using fluorescently-tagged proteins under the control of their endogenous promoters (Huang et al., 2009; Sabino et al., 2011; Kale et al., 2018). In *α-Cat-RNAi α-CatX* embryos, the levels of Ecad correlated with the strength of adhesion defects observed (Fig. 4A,C; for a summary of results see Fig. 4D), with lower levels of Ecad associated with more severe defects, as expected. Removal of M2 or M3 caused a significant reduction of cortical Ecad, suggesting that these domains predominantly contribute to the stability of CCC proteins at the junction (Fig 4A,C,D). This concurs with a reduction of junctional Arm in wing disc cells lacking M2 or M3 (Sarpal et al., 2019). In contrast, the M1 domain has an inhibitory effect on Ecad. *α-Cat-RNAi α-CatR-ΔM1* embryos showed elevated cortical enrichment of Ecad (Fig. 4A,C,D), consistent with an increase in junctional Arm in α-CatR-ΔM1 expressing wing disc epithelium (Sarpal et al., 2019; Alégot et al., 2019). An increase in Ecad levels as a result of the absence of M1 was also seen when comparing α-Cat-ΔM23 and α-Cat-ΔM embryos, and α-Cat-ΔM appears to be better maintained at the membrane and is less punctate than α-Cat-ΔM23 (Figs. 4 and S2A). We conclude that the M region regulates junctional stability with M1 and M2/3 having opposing effects. Whereas M2/3 stabilizes the junction, M1 limits junctional Ecad.

**Figure 4.**
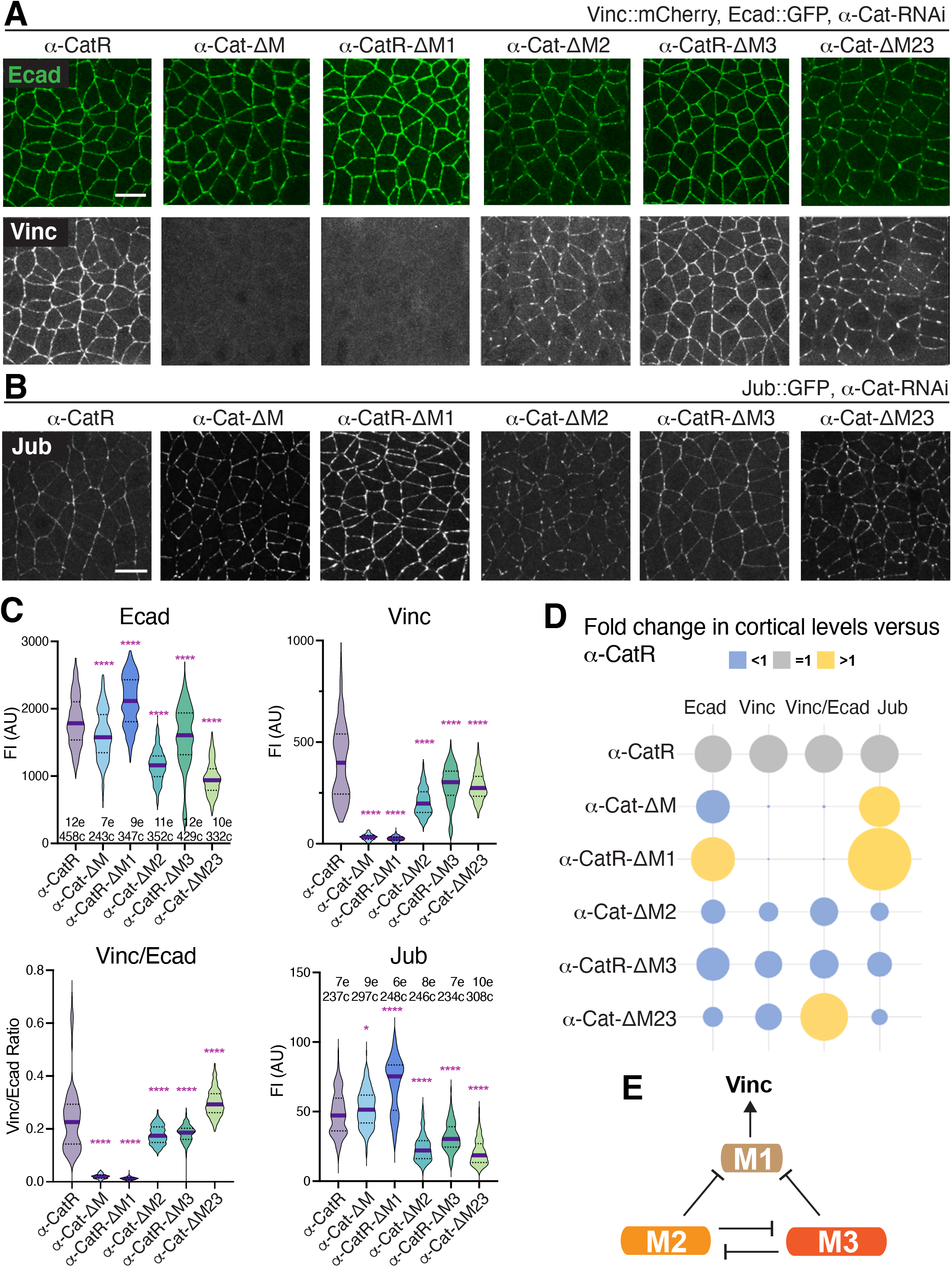
The M region regulates the levels of AJ components. **(A)** Stills of live *α-Cat-RNAi α-CatX* embryos at stage 8 expressing Ecad::GFP and Vinc::mCherry. **(B)** Stills of live *α-Cat-RNAi α-CatX* embryos at stage 8 expressing Jub::GFP. Scale bar for (A) and (B), 10μm. **(C)** Fluorescent intensities (FI) of junctional signal of Vinc::mCherry, Ecad::GFP, and Jub::GFP in *α-Cat-RNAi α-CatX* embryos at stage 8. N= number of cells (c), and embryos (e). The ratio of cortical Vinc to Ecad per cell is given as Vinc/Ecad. Significance calculated by ordinary one-way ANOVA (**** = P <0.0001, * =P <0.0332). **(D)** Balloon plot summarizing fold changes of junctional fluorescent signals. **(E)** Schematic illustration of the functional relationship between M1, M2, and M3 in respect to the recruitment of Vinc to the AJ.

### M2 and M3 negatively regulate M1 function

M1 is required for all detectable junctional Vinc signal in gastrulating embryos (Fig. 4A,C,D). As removal of M2 and M3 also reduces Ecad, we wanted to know whether the reduction in Vinc in these conditions is explained by a reduction of the CCC. We therefore calculated the ratio of cortical Vinc to Ecad per cell (expressed within the same embryo). We found that the M2 and M3 domains contribute to Vinc junctional levels even when normalized to the amount of remaining Ecad (Fig. 4A,C,D). Removal of both M2 and M3 together significantly increases the Vinc/Ecad ratio compared to α-CatR control (Fig. 4A,C,D). Together, this suggests that both M2 and M3 individually inhibit M1, but also each other, such that removing M2 and M3 together relieves all inhibition of Vinc recruitment (Fig. 4E). Removing M2 alone leaves M3 free to inhibit the M1-Vinc interaction and vice versa. This is in agreement with work using cells in culture identifying a masking effect between domains (Yonemura et al., 2010; Matsuzawa et al., 2018; Sakakibara et al., 2020), and provides evidence within an animal model to support the existence of autoinhibition among M region domains.

### M1 inhibits the α-Cat-Jub interaction during germband extension

In wing disc epithelial cells expressing M2 or M3 deletions instead of endogenous α-Cat, Jub is reduced likely due to an overall reduction in CCC levels. However, Jub levels are strikingly increased in cells lacking M1 (Sarpal et al., 2019; Alégot et al., 2019). We observed a similar pattern in the early embryo. Embryos where M1 is removed showed a dramatic increase in Jub at the junctions (Fig. 4B,C,D). In contrast, Jub levels are reduced in *α-Cat-RNAi* embryos expressing α-Cat-ΔM2, α-CatR-ΔM3, or α-Cat-ΔM23, suggesting that the M2 and M3 domains support junctional Jub. Despite lacking M2 and M3, embryos expressing α-Cat*Δ*M experience an enrichment of Jub (Fig. 4B,C,D). Thus, as shown for the wing disc epithelium (Sarpal et al., 2019; Alégot et al., 2019), M1 also acts as an inhibitor of junctional Jub recruitment in the embryonic ectoderm. Ectopic exposure of the M1 domain in *α-Cat-RNAi α-Cat-ΔM23* embryos produces the largest reduction in Ecad, Jub (Fig. 4), and the strongest adhesion defects (Figs. 1C, 2, S2F). As junctional Jub has a positive role in cell adhesion in the ectoderm (Razzell et al., 2018), we conclude that M1 is a negative regulator of cell adhesion, a role normally limited by the functions of M2 and M3, an inhibition resolved by the force-dependent unfurling of the M region.

### The M2 domain is required for the mechanosensitive enrichment of Jub at high tension edges

We also found that manipulation of the M region significantly affected the mechanosensitive recruitment of Jub. Jub is recruited to the membrane in response to cytoskeletal tension (Rauskolb et al., 2019; Razzell et al., 2018; Alégot et al., 2019), In wing disc epithelium, removal of the M1 domain causes Jub to become hyper-recruited to junctions without an increase in tissue tension (Sarpal et al., 2019; Alégot et al., 2019). How this occurs is unclear as the Jub-binding site is not within the α-Cat M region, but in the N2 domain (Marie et al., 2003; Alégot et al., 2019; Sarpal et al., 2019). A requirement for the α-Cat N-terminal domain for the junctional recruitment of Jub was confirmed in the embryo, as rescue with a construct which lacks this domain (DEcadΔβ::α-Cat-ABD) depleted Jub levels to the same degree as α-Cat knockdown (Fig. 5A-C). During germband extension, where vertical edges are under higher tension than horizontal contacts, this leads to the planar polarized enrichment of Jub to vertical edges (Razzell et al., 2018). This provided us with a system in which we could test Jub response to tension. We determined the planar polarity of Jub in *α-Cat-RNAi α-CatX* embryos by measuring the average fluorescent intensity of Jub along vertical edges divided by that of horizontal edges within the same embryo. This analysis revealed that the M2 domain is in fact required for the specific enrichment of Jub to vertical cell contacts (Fig. 5D). Expression of α-Cat-ΔM, α-Cat-ΔM23, and α-Cat-ΔM2 failed to rescue the enrichment of Jub to edges approaching 90° (where the anterior-posterior axis = 0°), and in some instances a reversal of Jub planar polarity was seen with higher Jub levels at horizontal versus vertical edges (Figs. 5D and S3A).

**Figure 5.**
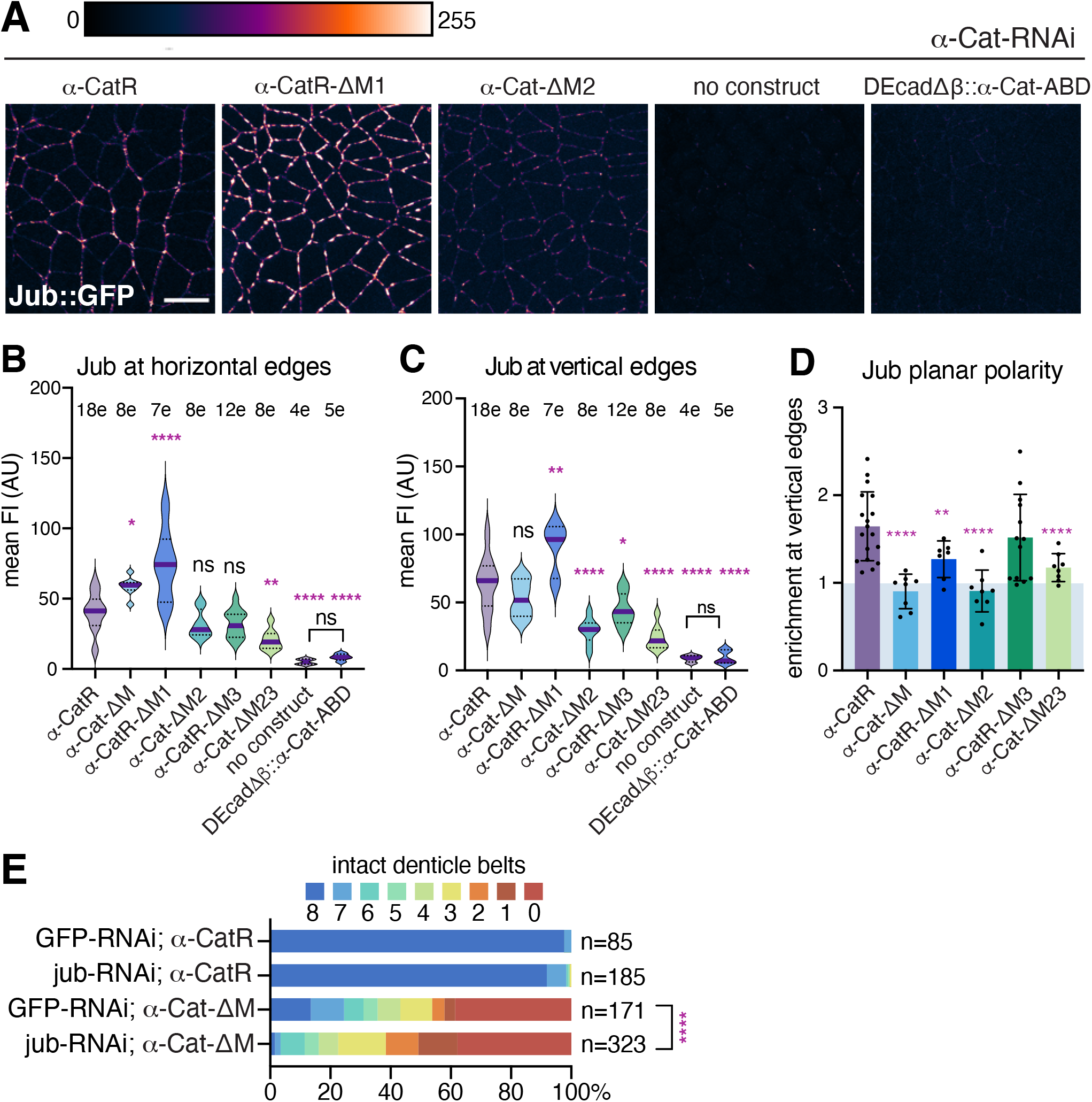
M region supports mechanosensitive recruitment of Jub to AJs. **(A)** Stills from live *α-Cat-RNAi α-CatX* embryos expressing Jub::GFP. Scale bar, 10μm. **(B,C)** The average fluorescent intensity (FI) of Jub (with background cytoplasmic signal subtracted) for *α-Cat-RNAi α-CatX* embryos at stage 8 is plotted for (B) horizontal edges (0-15°) and (C) vertical edges (75-90°, 0=anterior-posterior axis). N= number of embryos (e). **(D)** Quantification of planar polarity of Jub is plotted per embryo as average FI of vertical edges divided by that of horizontal edges. The ratio in (D) is derived from the data in (B) and (C). For (B) and (C), significance is calculated by ordinary one-way ANOVA, and for (D) by two-way ANOVA (**** = P <0.0001, ***= P <0.0002, **=P<0.0021, *=P <0.0332). See Fig S3 for more detailed breakdown of enrichment of fluorescent intensity by angles of edges. **(E)** Analysis of cuticle defects of *α-Cat-RNAi α-CatX* embryos and either *GFP-RNAi* as a control or *Jub-RNAi*. Chi-square test (**** = P <0.0001).

The planar polarity of Jub in *α-Cat-RNAi α-Cat-ΔM1* embryos is also significantly reduced, but in this case this is due to a constitutive enrichment of Jub to all cell edges, regardless of the state of tension (Figs. 5A-D and S3A). Similarly, in wing disc epithelium, loss of M1 desensitizes junctional Jub recruitment to a reduction of myosin activity (Alégot et al., 2019). The planar polarity of Jub thus requires M2 for the mechanosensitive recruitment to higher tension edges, while M1 limits Jub at lower tension edges. Although Jub is required for normal germband extension, its planar polarity is not essential for adhesion or germband extension (Razzell et al., 2018). The M2 domain may therefore function by supporting the necessary amount of Jub at higher tension edges, and hence support Ecad membrane stability against enhanced actomyosin contractility. As junctional Jub is more enriched in α-Cat-ΔM than α-Cat-ΔM23 embryos, we wondered whether the lack of M1-mediated inhibition of Jub helps to explain the difference in rescue of cell adhesion by these two constructs. Indeed, depletion of Jub by RNAi worsened epithelial integrity of *α-Cat-RNAi α-Cat-ΔM* embryos (Fig. 5E). These results suggest that Jub and the M2 domain cooperate with each other to support AJs under morphogenetic stress, and underline a requirement for the M region for the mechanosensitive recruitment of Jub to edges under higher tension.

### The M region reinforces Ecad against enhanced actomyosin contraction

In contrast to the enrichment of the α-Cat binding partners Jub, Vinc and Cno at vertical, high tension edges during germband extension (Blankenship et al., 2006; Sawyer et al., 2011; Kale et al., 2018; Razzell et al., 2018), AJ components such as Baz, Arm, α-Cat and Ecad become planar polarized and enriched at horizontal edges (Paré and Zallen, 2020). Within an Ecad::GFP, GAP43::mCherry background, embryos expressing *α-Cat-RNAi α-Cat-ΔM* and *α-Cat-RNAi α-Cat-ΔM23* showed significantly enhanced planar polarized localization of Ecad. Ecad was reduced specifically from the actomyosin-enriched vertical edges compared to α-CatR controls (Fig. 6A), suggesting that the M2 and M3 domains support Ecad at high-tension edges. Interestingly, within an Ecad::GFP, Vinc::mCherry background, a reversal of Ecad planar polarity was seen in *α-Cat-RNAi α-Cat-ΔM23* embryos (in which Vinc is ectopically enriched at AJs), while there was no change to Ecad planar polarity in embryos rescued by α-Cat-ΔM or α-CatR-ΔM1 (which cannot bind Vinc) (Fig. 6A). Expression of Vinc::mCherry to assess Vinc distribution provided an additional genomic copy of Vinc (Kale et al., 2018), representing an over-expression condition. This additional copy of Vinc also improves adhesion in the lateral ectoderm of *α-Cat-RNAi α-Cat-ΔM23* embryos compared to controls without Vinc::mCherry (Fig. 6B). As another method to enhance Vinc activity, we expressed the constitutively active Vinc-CO mutation (Maartens et al., 2016). This caused a strong enrichment of Ecad (Fig. 6C), and a reversal of Ecad planar polarity such that Ecad is enriched at vertical edges (Fig. 6D). These results suggest that the M1-Vinc interaction can reinforce Ecad recruitment or stability in response to actomyosin contraction and, consequently, cell adhesion during Drosophila gastrulation. However, as the M1 domain is dispensable for adhesion and embryonic development, it is unlikely that the recruitment of Vinc is the primary mode of junctional reinforcement through the M region in wildtype. Instead, our data suggests that M1 functions as an inhibitor of AJs.

**Figure 6.**
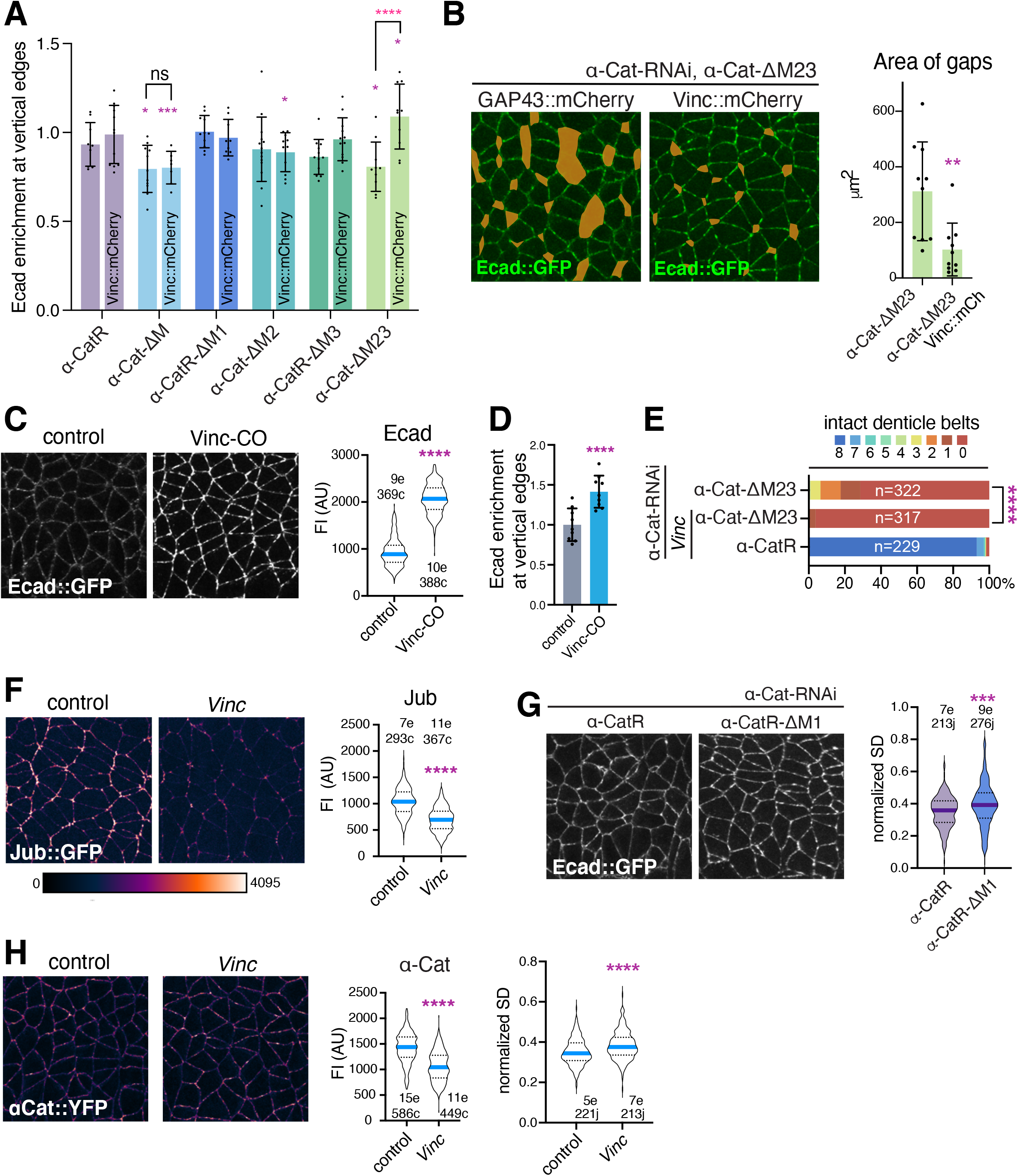
The M1-Vinc interaction supports Ecad stability and reinforces adhesion. **(A)** Ecad planar polarity (enrichment at vertical edges) is shown in *α-Cat-RNAi α-CatX* embryos at stage 8 expressing either GAP43::mCherry or Vinc::mCherry. **(B)** An extra copy of Vinc (Vinc::mCherry) improves adhesion in the lateral ectoderm in *α-Cat-RNAi α-Cat-ΔM23* embryos at stage 8. **(C, D)** Comparison of control and Vinc-CO embryos expressing Ecad::GFP. Stills from live embryos at stage 8. Fluorescent intensity (FI) of Ecad and Ecad planar polarity (D). **(E)** Denticle belt count of *α-Cat-RNAi α-Cat-ΔM23* embryos with and without a *Vinc* maternal-zygotic (MZ) null mutant background. **(F)** Live imaged stills comparing Jub::GFP signal in wildtype and a *Vinc* MZ null mutant embryo at stage 8. **(G)** *α-Cat-RNAi α-CatR-ΔM1* embryos at stage 8 show less uniform junctional distribution of Ecad::GFP compared to *α-Cat-RNAi α-CatR* controls as indicated by the increased normalized standard deviation (SD) of Ecad::GFP signal at AJs. **(H)** Junctional α-Cat::YFP signal (same color map in F), in stage 8 embryos is reduced and less uniformly distributed in the absence of *Vinc,* as indicated by an increase of the normalized SD of α-Cat::YFP signal. N= junctions (j) and embryos (e) in (G) and (I). N= cells (c) and embryos (e) in (E) and (K). Significance is calculated by two way ANOVA for (A), Chi-square for (E), and Mann-Whitney two-tailed test for (B), (C), (D), (F), (G), and (H) (**** = P <0.0001, ***= P <0.0002, **=P<0.0021, *=P <0.0332).

### The M1 domain is required for the normal distribution of junctional Ecad through recruitment of Vinc

Since M1 has an inhibitory effect on Ecad and Jub, and over-exposure of M1 both enriches Vinc and causes more severe defects than removal of the whole M region, we wondered whether Vinc has an inhibitory effect on Ecad or Jub, and thus adhesion. A *Vinc* null mutant allele was incorporated into the rescue set-up to see if this would ameliorate epithelial integrity defects in *α-Cat-RNAi α-Cat-ΔM23* embryos. However, removal of Vinc in fact worsened the defects in α-Cat-ΔM23 embryos, suggesting that Vinc cooperates with the M2 and M3 domains to support adhesion (Fig 6E). Furthermore, we confirmed that *Vinc* null mutants have no discernable epithelial defects during gastrulation, as previously reported (Alatortsev et al., 1997; Maartens et al., 2016), but found subtly yet significantly reduced α-Cat and Jub levels (Fig. 6F,H). Although M1 is similarly dispensable for embryonic development, abnormally large clusters of Ecad, Jub, and α-Cat-ΔM1 were seen in *α-Cat-RNAi α-Cat-ΔM1* embryos along the junctions compared to α-CatR expressing controls (Figs. 5A, 6G, S4A). We used the normalized standard deviation of Ecad to estimate the fragmentation of junctional signal, and this was significantly increased in the absence of M1 (Fig. 6G). Likewise, junctional α-Cat signal is more fragmented in *Vinc* embryos (Fig. 6H). M1 recruitment of Vinc therefore does support a more uniform distribution of the CCC at AJs, but the impact of its loss has no apparent phenotypic consequences. Our observations suggest that the inhibitory effect the M1 domain has on adhesion is not due to Vinc recruitment, and that the M1 domain plays a dual inhibitory and supportive role at AJs.

### Cno requires M2/M3 for its mechanosensitive enrichment to TCJs, but is cortically recruited independently of the α-Cat M region

Loss of either Vinc or Jub alone did not produce significant adhesion defects in Drosophila embryos (Maartens et al., 2016; Razzell et al., 2018), and also Jub knockdown in a *Vinc* mutant background did not enhance embryonic lethality. However, loss of Vinc or Jub enhanced epithelial defects in M region deletion construct expressing embryos (Figs. 5E and 6E), raising the possibility that they work in parallel with an additional interaction partner of the M region, contributing to its mechanosensory output. As a known binding partner of the mammalian α-catenin M2/M3 domains (Pokutta et al., 2002; Sakakibara et al., 2020), we examined therefore the Drosophila Afadin ortholog Cno to assess if force-dependent recruitment of Cno by the M region could help explain M region function in cell adhesion.

Although a masking effect of M1 on the M3-Afadin interaction was reported in MDCK cells (Sakakibara et al., 2020), in the Drosophila embryo we found that removal of either M3 or M1 led to an increase in junctional Cno (Fig. 7A,B). In fact, junctional Cno increased modestly with the expression of all M region deletions except for α-Cat-ΔM23, which showed a reduction but not a loss of Canoe at AJs (Fig. 7A,B). By stage 8, Cno is significantly reduced by the knockdown of α-Cat alone (Fig. 7A,B), when few cell junctions remain. Since Cno levels are increased in α-Cat-*Δ*M expressing embryos, the reduction of Cno in *α-Cat-RNAi α-Cat-ΔM23* embryos may be a consequence of the strong adhesion defects in this condition. Although the planar polarity of Cno in *α-Cat-RNAi α-CatR-ΔM3* embryos was reduced, Cno planar polarity was unaffected in *α-Cat-RNAi α-Cat-ΔM* (Fig. S3). This did not allow us to conclude that M3 is required for the enrichment of Cno at vertical edges.

**Figure 7.**
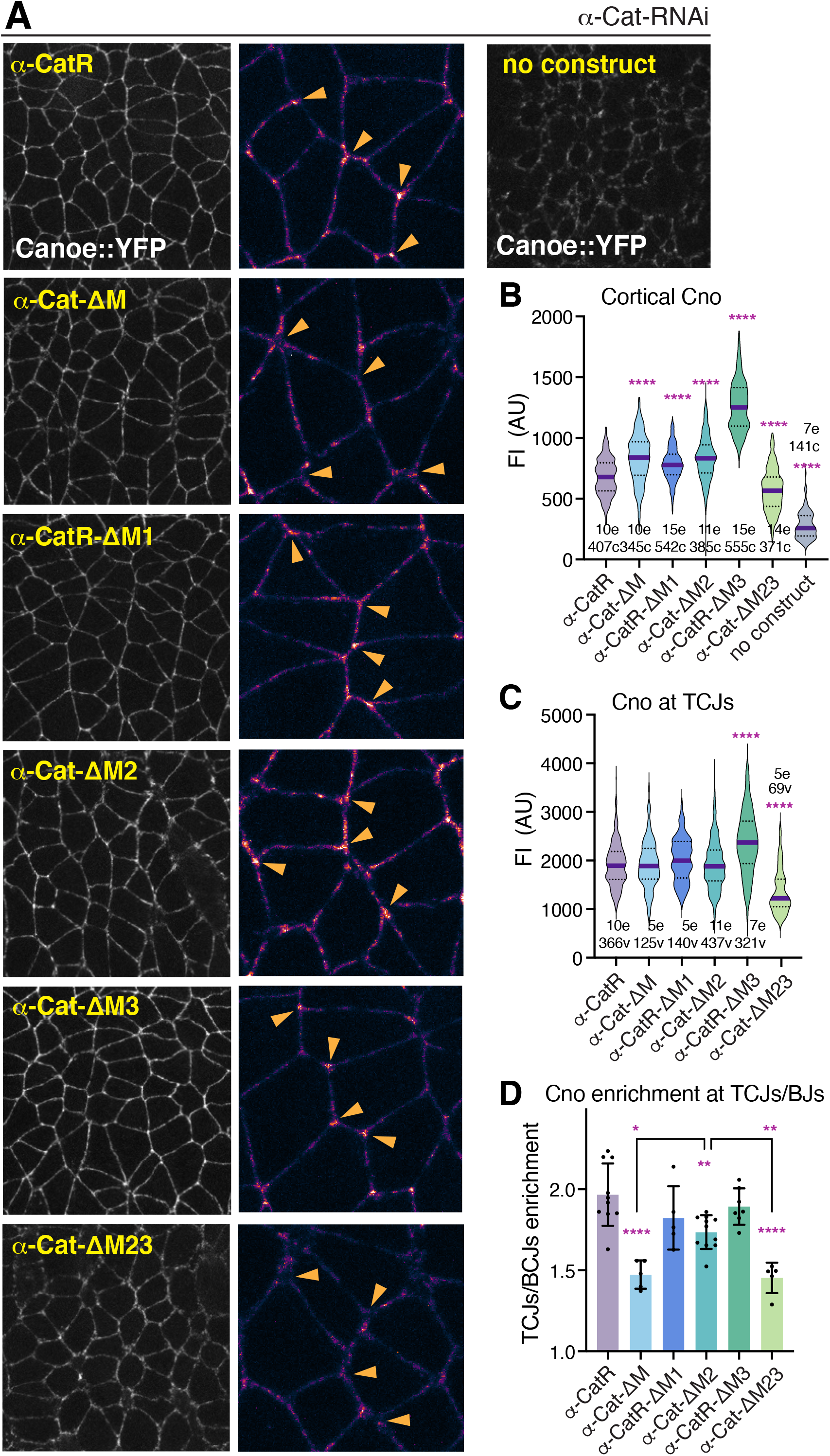
Mechanosensitive recruitment of Cno to TCJs requires the α-Cat M region. **(A)** Stills of lateral ectoderm at stage 8 of *α-Cat-RNAi α-CatX* live embryos expressing Cno::YFP. Orange arrowheads in close-ups at right point to TJCs. **(B)** Cortical levels of Cno of *α-Cat-RNAi α-CatX* embryos. N=number of embryos (e) and cells (c). **(C)** Levels of Cno measured at intact TCJs in *α-Cat-RNAi α-CatX* embryos. N=number of embryos (e) and vertices (v). **(D)** Ratio of the mean fluorescence of Cno at TCJs versus bicellular junctions (BCJs) is plotted per embryo. Significance is given by one-way ANOVA (**** = P <0.0001, ***= P <0.0002, **=P<0.0021, *=P <0.0332).

Cno is normally elevated both at vertical BCJs during gastrulation and also at TCJs (Sawyer et al., 2009; 2011). The enrichment of Cno at TCJs is responsive to cytoskeletal tension (Yu and Zallen, 2020). We therefore assessed Cno levels at vertices as a TCJ/BCJ ratio within the same embryo. Although the minimal Afadin binding site of αE-catenin is located within M3 (Pokutta et al., 2002; Sakakibara et al., 2020), removing M3 alone had no effect on the TCJ enrichment of Cno (Fig 7A,D). However, the enrichment of Cno at vertices was significantly reduced in embryos lacking M2, and this is worsened when M3 is also removed (Fig. 7A,D). These results suggest that M2 and M3 can interact with Cno independently and that M3 supports M2 in the mechanosensitive recruitment of Cno to TCJs.

The pools of Cno at BCJs versus TCJs thus appeared to have different requirements for the M region. Interestingly, Cno at BCJs does not show a response to experimental changes in tension (Yu and Zallen, 2020). Measurements of cortical Cno represent predominantly the bicellular pool, which does not require the M region (Fig. 7B). Loss of the M region also does not abolish Cno localization to TCJs (Fig. 7C,D). Cno must therefore have alternate pathways to associate with the plasma membrane. During cellularization, Cno recruitment is unaffected by the loss of Arm, and instead requires the small GTPase Rap1 (Sawyer et al., 2009). Echinoid (Ed) is thought to cooperate with Ecad to support Cno recruitment (Wei et al., 2005; Sawyer et al., 2009). However, even in a naturally occurring Ed deficient tissue, the amnioserosa (Lin et al., 2017), junctional recruitment of Cno was not depleted in *α-Cat-RNAi α-CatR-ΔM3* or *α-Cat-RNAi α-CatR-ΔM* embryos (Fig. S4). Deletion of M1 exposes the Cno binding site, but bicellular α-Cat-ΔM1 signal does not colocalize better with Cno than α-CatR control when expressed in *α-Cat-RNAi* embryos (Fig. S4), further supporting the conclusion that Cno can localize to BCJs independently of the M region. Finally, α-Cat constructs were also consistently located more basally than Cno at stage 8 (Fig. S4). Taken together, these findings suggest that Cno is recruited to the membrane independently of the α-Cat M region, but that the mechanosensitive recruitment of a pool of Cno to TCJs requires an interaction with the α-Cat M2 and M3 domains.

### The α-Cat M region acts in parallel to Cno to support cell adhesion

If M region mechanosensing supports adhesion by recruiting Cno as a downstream effector, then compromising Cno within an embryo lacking the M2 and/or M3 domains should not worsen adhesion defects. However, Cno knockdown enhanced the epithelial defects of all α-Cat deletion constructs (Fig. 8A). Furthermore, overexpression of Cno improves epithelial integrity in embryos expressing *α-Cat-RNAi* alone (Fig. 8B), and constructs lacking the M2 and M3 domain (Fig. 8C). In particular, overexpression of Cno significantly ameliorated the loss of ventral epithelium in *α-Cat-RNAi α-Cat-ΔM23* and *α-Cat-ΔM* embryos, which are predicted to abolish Cno binding, as well as α-Cat-*Δ*M2 (Fig. 8C,D). Together with the M region-independent association of Cno with AJs, these observations suggest that Cno supports adhesion in an α-Cat-independent parallel pathway. In further support of this conclusion we found that knockdown of the Rap1 GTPase, which contributes to Cno membrane recruitment and activation (Sawyer et al., 2011; Bonello et al., 2018; Perez-Vale et al., 2021), enhances the defects in *α-Cat-RNAi α-CatR-ΔM* embryos (Fig. 8E). These results suggest that Rap1 interaction with Cno shares some parallel function with the M region to support adhesion. Although these results do not rule out a role for M region-dependent Cno recruitment, it argues against Cno acting solely as an effector of α-Cat mechanosensing. Instead, Cno appears to have a redundant role to the M region, and to the M1 domain in particular as knockdown of Cno enhances loss of junctional Ecad and cell adhesion defects in α-CatR-ΔM1 embryos compared to α-CatR controls (Fig. 8A,D).

**Figure 8.**
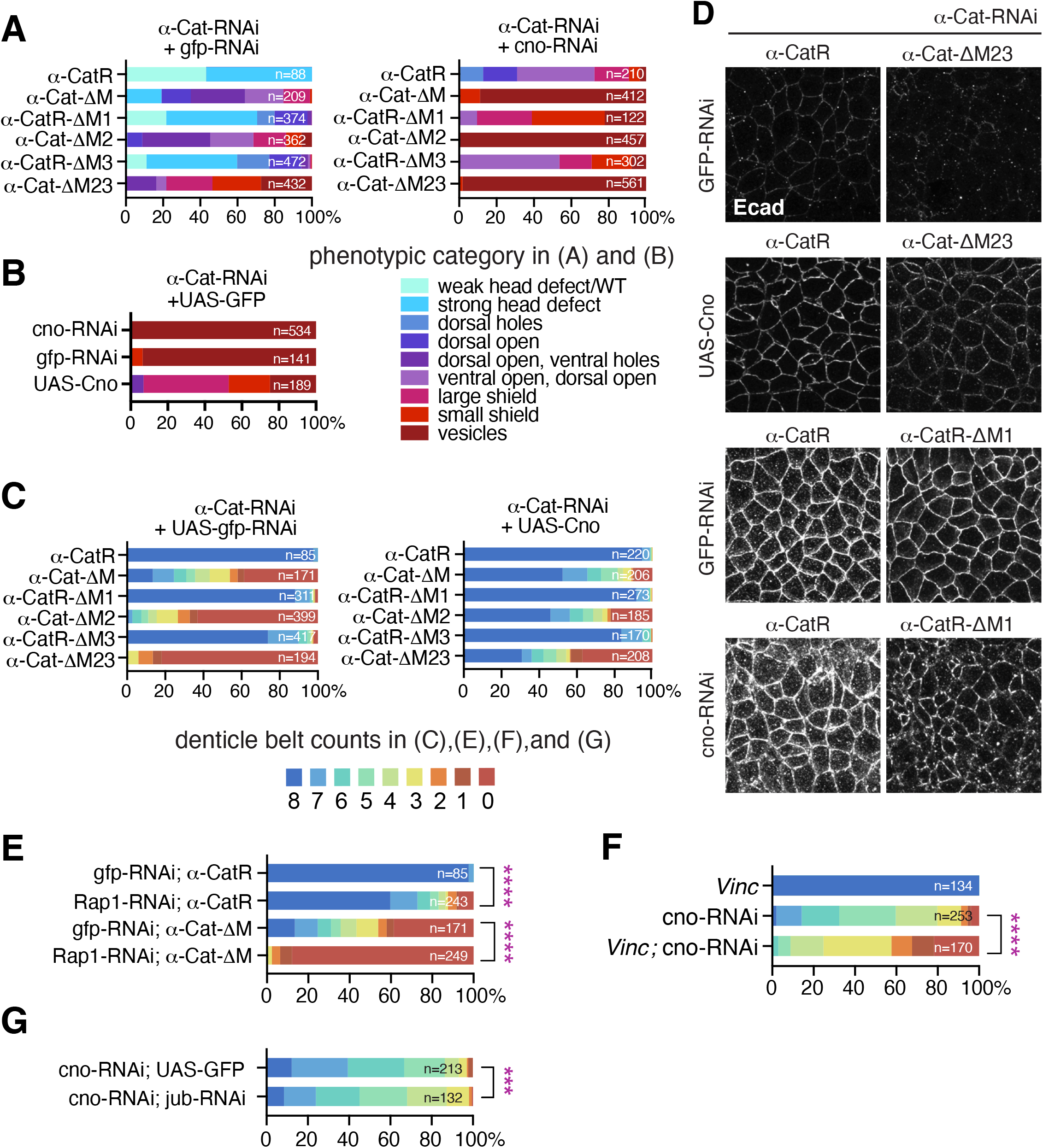
Cno supports Ecad stability in a parallel pathway to the α-Cat M region. **(A)** Quantification of cuticle defects from c*no-RNAi α-Cat-RNAi α-CatX* and *gfp-RNAi α-Cat-RNAi α-CatX* embryos. Phenotypic categories are color-coded and examples are given in Fig. S5. **(B)** Overexpression of Cno improves epithelial integrity in *α-Cat-RNAi* embryos. **(C)** Denticle belt count for *α-Cat-RNAi α-CatX* embryos overexpressing Cno or *GFP-RNAi* as control. *gfp-RNAi* controls same as in Fig. 5E. **(D)** Ectoderm at stage 8/9 of *α-Cat-RNAi α-CatX* embryos expressing either *UAS-cno*, *cno-RNAi* or *UAS-gfp-RNAi* as control immunostained for Ecad. Ecad levels are enhanced by Cno overexpression, ameliorating defects found in α-Cat-ΔM23 embryos, whereas Cno knockdown causes an enhancement of AJ fragmentation in α-CatR-ΔM1 embryos compared to control. **(E)** *Rap1-RNAi* enhances defects in *α-Cat-RNAi α-CatR* and *α-Cat-RNAi α-Cat-ΔM* embryos.*gFP-RNAi* controls same as in (C). Chi-square test, ****=P <0.0001. **(F)** *Vinc* null mutant enhances defects caused by *cno-RNAi*. Chi-square test, ****=P <0.0001. **(G)** *jub-RNAi* enhances defects caused by *cno-RNAi*. Chi-square test, ***= P <0.0002.

### α-Cat binding partners and Cno form a cooperative network to support adhesion

We wondered whether the recruitment of Vinc by M1 is redundant with Cno function. Epithelial integrity was significantly worsened when Vinc is removed from a *cno-RNAi* background, consistent with functional redundancy between Cno and the M1-Vinc interaction (Fig. 8F). Functional redundancy between Cno and the M1 domain could explain why M1, and similarly, Vinc are dispensable in the embryo. Furthermore, double knockdown of Jub and Cno significantly enhanced adhesion defects compared to Cno knockdown alone (Fig. 8G). Such findings argue for a model where multiple ways to stabilize Ecad build a robust partially redundant network of interactions to foster AJ stability. Our evidence supports a model where the α-Cat M region itself, and the α-Cat binding partners Vinc and Jub cooperate with Cno to promote cell adhesion.

## Discussion

α-catenin is essential for cell adhesion in most animal tissues. α-catenin acts as a physical linker between the cadherin/β-catenin complex and the actin cytoskeleton (Desai et al., 2013; Buckley et al., 2014), and through its mechanosensory properties has the potential to modify interactions between cadherins and the cytoskeleton in response to external or internal forces (Yonemura et al., 2010; Le Duc et al., 2010; Borghi et al., 2012; Buckley et al., 2014; Ishiyama et al., 2018; Xu et al., 2020). Two mechanosensory regions within α-catenin have been identified: the C-terminal actin-binding domain of α-catenin increases binding strength to F-actin as force is applied (Buckley et al., 2014; Ishiyama et al., 2018; Xu et al., 2020), and the central M region of α-catenin that acts principally by modifying interactions between α-catenin and binding partners in response to force (Yonemura et al., 2010; Ishiyama et al., 2013; Thomas et al., 2013; Yao et al., 2014; Li et al., 2015; Maki et al., 2016; 2018; Seddiki et al., 2017; Barrick et al., 2018; Matsuzawa et al., 2018; Terekhova et al., 2019; Sarpal et al., 2019; Alegot et al., 2019). How α-catenin mechanosensing cooperates with multiple known α-catenin binding partners that can also bind to F-actin to dynamically adjust adhesion during tissue morphogenesis has remained largely unexplored.

We determined that the M region of Drosophila α-Cat plays a key role in maintaining cell adhesion during mesoderm invagination and cell intercalation that drives axis elongation in the early embryo. Taking advantage of the well-described tension landscape in the ectoderm as it undergoes cell intercalation during germband extension (Bertet et al., 2004; Fernandez-Gonzalez et al., 2009; Tetley et al., 2016), we were able to identify three different tension states of α-Cat that correspond to the differential recruitment of binding partners (Fig. 9). (i) Moderate amounts of Vinc and Jub are recruited to AJs at low-tension BCJs (=horizontal junctions in the extending germband); (ii) At medium-tension BCJs (=vertical junctions in the extending germband) recruitment of Vinc and Jub is significantly increased; and (iii) at TCJs, which experience the highest levels of tension, Vinc and Jub are further elevated, and Cno is enriched in response to α-Cat mechanosensing. Recruitment of Vinc and Jub to AJs critically depends on physical interaction between Vinc and the M1 domain and Jub and the N2 domain. In contrast, Cno localizes at AJs via mechanisms that are independent of α-Cat and supports cell adhesion in a parallel pathway. Only the mechanosensitive enrichment of Cno at tricellular vertices (Yu and Zallen, 2020) requires its interaction with the M2 and M3 domains of α-Cat.

**Figure 9.**
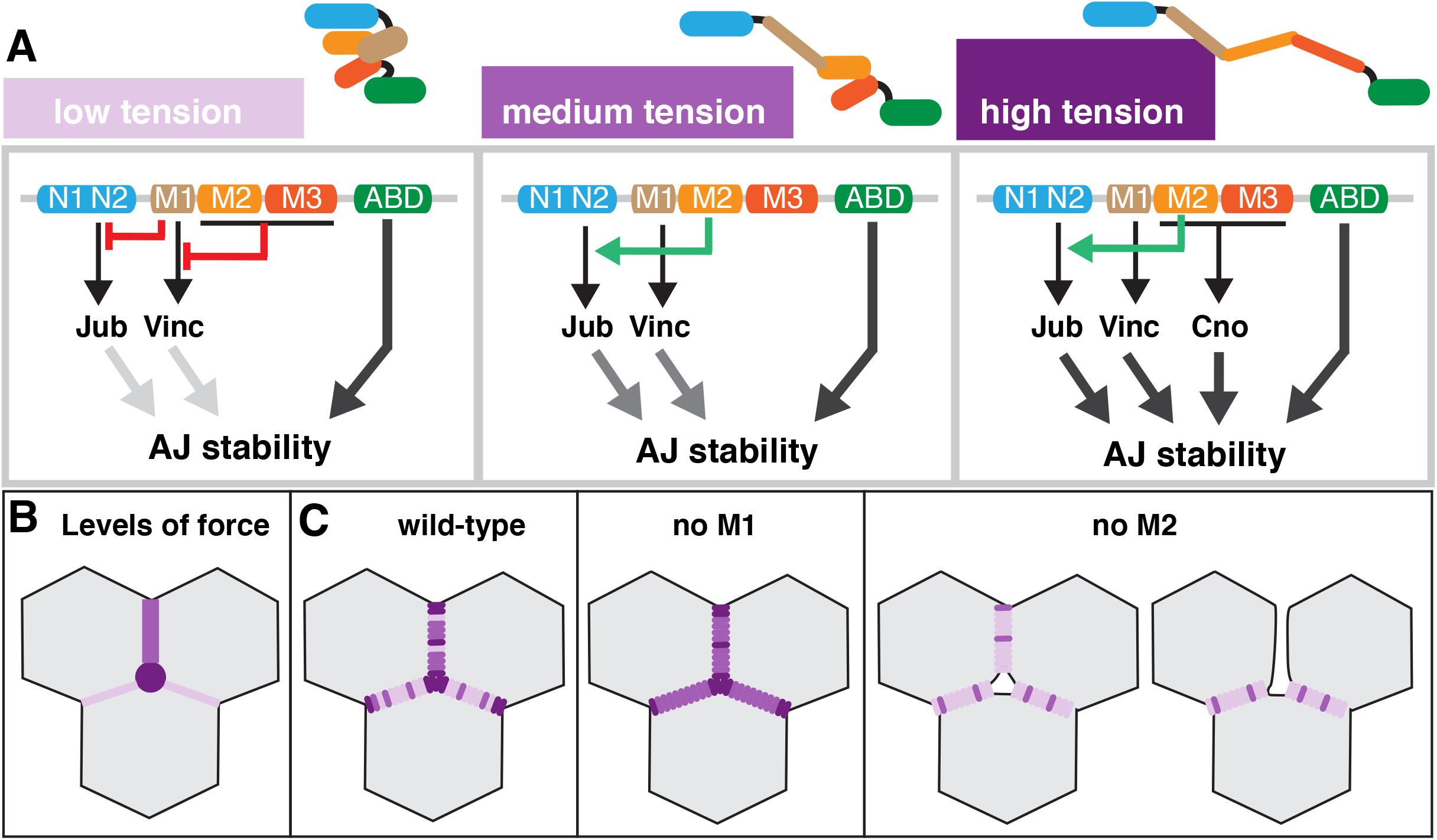
Model of α-Cat M region mechanosensing in the Drosophila ectoderm during germband extension. **(A)** Schematic illustrating three distinct conformation of the α-Cat M-region found at low, medium and high tension cell contacts. demonstrating the distinct contributions of M-region subdomains to the recruitment of binding partners. M-region subdomains have an inhibitory effect on each other. **(B)** Diagram illustrating the tension landscape in the ectoderm during germband extension, showing low tension contacts (horizontal edges), medium tension contacts (vertical edges) and high tension contacts (TCJs/vertices). **(C)** Schematic illustration of anticipated distribution of the three α-Cat conformation highlighted in (A) in wild-type, *α-Cat-RNAi α-CatR-ΔM1* embryos and *α-Cat-RNAi α-Cat-ΔM2*. See text for further discussion.

The M region of α-catenin is stabilized by a dynamic network of salt-bridges (Ishiyama et al., 2013; Li et al., 2015). Single molecule force measurements (Yao et al., 2014; Maki et al., 2016) suggest that the autoinhibited conformation can open when forces above approximately 5 pN, the range of force generated by a single myosin motor protein (Finer et al., 1994). A 5 pN force unfurls the M1 domain which becomes accessible to Vinc, an interaction that stabilizes the open (M1 unfurled) conformation of α-catenin (Yao et al., 2014; Maki et al., 2016). A further conformational change, likely affecting the M2 and M3 domains takes place when a ∼10-15 pN force is applied (Yao et al., 2014). Finally, the M1-Vinc interaction is destabilized at forces above 30 pN, and Vinc dissociates from α-catenin (Yao et al., 2014). Projecting these findings onto our results, we suggest that the difference between horizontal and vertical edges is in the frequency by which α-Cat is exposed to low (≦ 5 pN) or medium (5-10 pN) forces (Fig. 9 A-C). At vertical edges, medium tension/forces prevail that cause the majority of α-Cat molecules to adopt an M1 unfurled conformation which allows both Vinc binding to M1 and Jub recruitment to N2. The unfurled M1 conformation represents a smaller fraction of α-Cat molecules at horizontal edges causing a reduced amount of Vinc and Jub recruitment. Finally, forces between ∼10 and 30 pN per individual α-Cat molecule at TCJs would allow α-Cat to engage not only high levels of Vinc and Jub but also interact with Cno to further stabilize AJs (Fig. 9 A-C). A majority of α-Cat molecules are unlikely to experience forces above 30 pN as a loss of Vinc from the membrane is not observed even at TCJs.

The M1, M2 and M3 domains of α-Cat make distinct contributions to cell adhesion. M2 has a major role in supporting surface stability of the CCC and is required for the mechanosensitive AJ enrichment of Jub, which further stabilizes the junction. Together with M3, M2 supports Cno recruitment to TCJs to further reinforce AJs (Fig. 9). Recently, mechanical coupling between M2 and N2 domains has been identified (Terekhova et al., 2019), as well as direct, transient interaction between the disordered N-terminal region of *β*-catenin and the M region (Bush et al., 2019), suggesting that M2 may participate in strengthening interactions at the N-terminal domain with Jub and Arm. In contrast, M1 negatively regulates adhesion by inhibiting Jub recruitment, this negative regulation is released by M1 unfurling in response to force. While M1 recruitment of Vinc supports AJs stability, this contribution is subtle and dispensable for adhesion in wild-type. A noticeable effect of the loss of M1 or Vinc was a somewhat enhanced clustering of the CCC suggesting that the M1-Vinc interaction supports the uniform distribution of the CCC at cell contacts. Genetic analysis suggests that the M1-Vinc interaction is redundant with Cno and that Cno and the M region act in parallel to support AJs.

The M23 and M1 domains appear to have opposing roles in adhesion, similar to their function in the regulation of tissue growth (Sarpal et al., 2019). Whereas M1 has an important role in growth regulation, it is largely dispensable for adhesion, and the opposite is largely true for the M2 and M3 domains. This suggests that M region mechanosensing acts similarly in cell adhesion and tissue growth and that the tissue-specific consequences are due to differences in the downstream pathways that impact on growth versus adhesion. Our data would also suggest that Vinc does not play a significant role in either function (Sarpal et al., 2019). However, our results imply that Vinc supports adhesion in a subtle and redundant way. This is consistent with a previously reported worsening of the zygotic *α-Cat* mutant phenotype in combination with a *Vinc* mutation (Jurado et al., 2016). In embryos lacking M2, cortical levels of both Jub and Vinc, as well as Cno enrichment at TCJs, are reduced. Effects on multiple binding partners at once could could explain the strong adhesion defects observed in these embryos. Additional α-Cat binding partners such as α-Actinin, which also interacts with M1 (Nieset et al., 1997), remain to be investigated in our system, and could contribute to the redundant, multivalent interactions at the CCC-actin interface.

Recent analysis of Cno function in the Drosophila ectoderm highlighted the need for multiple, partially redundant interactions between Cno, the plasma membrane, and the cytoskeleton in support of AJ stability during tissue morphogenesis (Manning et al., 2019; Perez-Vale et al., 2021). Thus, both Cno and α-Cat stabilize AJs through multiple interactions. It is unclear, however, whether Cno is a mechanosensor or only responds to mechanosensory inputs by α-Cat as we show here or by other mechanisms as Cno enrichment at TCJs is also enhanced by phosphorylation of Cno by the Abl non-receptor tyrosine kinase (Yu and Zallen, 2020). Whether phosphorylation of Cno modifies the interactions with α-Cat remains to be explored. Cno at TCJs needs to be dynamic to enable the normal vertex resolution during the cell rearrangements required for cell intercalation (Yu and Zallen, 2020).

Taken together, our work suggests that the α-Cat M region acts in cell adhesion to support embryonic morphogenesis through mechanosensitive modulation of interactions with Vinc, Jub, and Cno. The M1-Vinc interaction, dispensable on its own, shows redundancy with Cno function. While Vinc and Jub association with AJs is completely dependent on α-Cat mechanosensing, the association of Cno with AJs depend on α-Cat mechanosensing only for its enrichment at TCJs. Cno supports adhesion at bicellular contacts in a parallel pathway to α-Cat. We suggest that these redundant and cooperative multivalent interactions are the molecular basis of the mechanoresponsive dynamic stabilization of AJs that maintain tissue integrity as the tissue undergoes cell contacts changes.

## Materials and Methods

### Drosophila Genetics

Flies were raised on standard media at 25°C for all experiments. For rescue experiments, males carrying *mat-GAL4* (*P{matα4-GAL-VP16}67* (Häcker and Perrimon, 1998) and the α-Catenin constructs (inserted at *attP2*) were crossed to females carrying fluorescently tagged proteins of interest and *UAS-α-Cat-RNAi* (TRiP HMS00317, Transgenic RNAi project [TRiP]). F1 virgin females carrying mat-GAL4, fluorescent protein, and α-Cat construct were then crossed to OregonR wild-type males (except *jub::GFP* males were used for rescue experiments measuring *jub::GFP*) and the progeny of this cross were analyzed. Similarly, for double knockdown analysis, F1 virgin females carrying both *mat-GAL4* and transgenes were crossed to OregonR wild-type males and their progeny assessed. In *Vinc* mutant analysis, a complete genomic deletion of *Vinc, Vinc^102.1^* was used (Klapholz et al., 2015). See Supplemental Table 1 for details of crosses used for each figure. The following fly lines were employed:

### UAS constructs

*α-CatR*, *α-CatR-ΔM1*, *α-Cat-ΔM* (aka: *α-CatR-ΔM*), *α-CatR-ΔM3* (Sarpal et al., 2019); *α-Cat-ΔM2* (aka: *αCatΔVH2-N*), *α-Cat-ΔM23* (aka: *αCatΔVH2*), *DE-cadΔβ::α-CatABD* (aka: *DE-cadΔβ::VH3-CTD*); *UAS-GFP* (Desai et al., 2013); *Vinc-CO* (Maartens et al 2016); *UAS-Canoe* (aka: *CanoeFL::GFP*) (Bonello et al., 2018).

### RNAi lines

*α-Cat-RNAi* (TRiP HMS00317); *jub-RNAi* (TRiP HMS00714); *canoe-RNAi* (TRiP HMS00239 and GL00633); *Rap1-RNAi* (TRiP HMJ21898); were produced by the Transgenic RNAi Project at Harvard (Zirin et al., 2020). *GFP-RNAi* (BL41552, Perrimon, N.).

### Fluorescently tagged markers

*α-Cat::YFP* (α-CatCPTI002516); *Canoe::YFP* (cnoCPTI000590); and *Zipper::YFP* (zipCC01626) are derived from the CPTI protein trap project, Kyoto Stock Centre (Lye et al., 2014; Lowe et al., 2014). Other fluorescently tagged proteins are insertions under the control of the respective endogenous promoter: Ecad::GFP, aka: DEcad::GFP, *shg>DEcad::GFP* (Huang et al., 2009); *Vinc::mCherry* (recombined with Ecad::GFP) (Kale et al 2019); *jub::GFP* (Sabino et al., 2011) Fluorescently tagged proteins ubiquitously expressed in the early embryo: Ani-RBD aka:*Ubi>Anillin-RBD::GFP* (Munjal et al., 2015) and *sqh>GAP43::mCherry* (Martin et al., 2010).

### Preparation of cuticle and quantification of embryonic lethality

To analyze the terminal cuticle phenotype of embryos, eggs collected overnight at 25°C were aged for 2 days or 24 hours if viable, and then washed in water and dechorionated for 5 mins in 2.15% sodium hypochlorite. After a second wash, embryos were mounted on a slide into a 3:2 mixture of Hoyer’s medium and lactic acid and incubated at 85°C overnight. To determine the percentage of embryonic lethality, flies were allowed to lay eggs at 25°C for an 8 hour period, and between 100-300 eggs were counted and arranged into rows on a new agar plate. This plate was then examined 24 and 48 hours later to count hatched larvae and dead embryos.

### Antibody staining

*Drosophila* embryos were dechorionated in 2.15% sodium hypochlorite and then either heat-fixed in a salt solution (Miller et al., 1989) (Fig. 8, S4) or fixed for 20 min in 4% formaldehyde in a 1:1 PBS:heptane mixture (Fig S2). Primary antibodies used under heat fixation were anti-HA (rat monoclonal, 3F10; 1:500; Roche), anti-Canoe (rabbit 1:1000, a gift from Mark Peifer, University of North Carolina, Chapel Hill, NC, USA), anti-DE-cadherin (rat monoclonal DCAD2 1:25; Developmental Studies Hybridoma Bank). Under heptane fixation: anti-HA (mouse 16B12 1:100, Abcam).

### Analysis in follicular epithelium

A heat-shock inducible MARCM system was employed as previously described (Sarpal et al., 2012; Desai et al., 2013). For cell clones produced in the follicular epithelium, the percentage of rescued cells was calculated. The following recombinant lines were used for MARCM analysis: *hs–Flp FRT40A; da–Gal4 UAS–mCD8::GFP α*-*Cat*^1^*/TM6B*; *tub–Gal80 ubi*– α-*Cat FRT40A; act–Gal4 a-CatX α*-*Cat*^1^*/TM6b* (Sarpal et al., 2012; Desai et al., 2013; Sarpal et al., 2019).

### Imaging and signal intensity quantification

Live imaging was performed on dechorionated embryos mounted in Halocarbon Oil 25 between an oxygen permeable membrane and coverslip, and short time-lapse movies were acquired, similar to other previously described methods (Blankenship et al., 2006). Fluorescent images were acquired using a Leica TCS SP8 scanning confocal microscope on 40× or 63× objectives (HC PL APO CS2 with NAs of 1.30 and 1.40, respectively). A Carl Zeiss Axiophot2 microscope using a phase-contrast 20× lens (NA 0.5) connected to a Canon Rebel XSi camera was used to capture images of embryonic cuticles, and time-lapse movies were captured using this scope and a 10x lens (NA 0.14) to determine rates of germband extension. The number of embryos per genotype is listed on each figure, and in Supplemental Table 1.

Z-stacks were collected using a z-step size of 0.35-0.4 µm, starting from above the vitelline membrane and moving well through all visible apical junctional structures for 15-20 steps. Planes containing autofluorescence of the vitelline membrane were removed in post-processing and maximum projections were produced using Fiji (Schindelin et al., 2012). Adobe Illustrator was used to place confocal images and curves were adjusted in Adobe Photoshop only for Vinc::mCherry confocal images and images of cuticle preparations. For images shown using a colormap legend, the ‘gem’ lookup table was used. The same settings on confocal equipment and image processing were applied for all images within the same experiment.

### Analysis of early embryonic phenotypes and germband extension

*α-Cat-RNAi α-CatX* expressing embryos were live imaged during stages 6, 7 and 8 and categorization was determined for embryos within a Jub::GFP, Ecad::GFP, GAP43::mCh, or myosin::YFP background (Supplemental Table 1). For quantification of germband extension, the change in position of the proctodeal invagination after 40 minutes was normalized to the total length of the embryo.

### Fluorescent Intensity

Analysis of cortical levels of fluorescence was performed using the Matlab script SIESTA (scientific image segmentation and analysis (Fernandez-Gonzalez and Zallen, 2011). For a number of embryos (e), a polyline 3 pixels in width was drawn along the perimeter of each cell (c), and the mean fluorescent intensity along this line, minus the mean fluorescent intensity within the center of the cell, was plotted as cortical fluorescent intensity (FI). Total junctional myosin is given as the sum of the mean fluorescence along the perimeter and mean fluorescence within the perimeter for each cell (total FI). Junctional/total myosin fraction is given as the average FI of myosin at the cell perimeter (junctional) out of total myosin (fraction of total FI). For comparison of Ani-RBD at the edge of gaps versus cells not in contact with a gap, the average fluorescent intensity along the perimeter (perimeter FI) was plotted for polylines drawn along cells (intact) versus along the gap (gap).

### Planar polarity

Using SIESTA, trajectories 3 pixels in width were drawn along individual cell edges, avoiding overlap or vertices and these edges were grouped into 15° bins, reflected about the DV axis. For each bin, the cytoplasmic fluorescence of the image was subtracted from the average fluorescent intensity of cell edges. This value was then divided by that of the 0-15 (horizontal) bin. Enrichment to vertical edges is plotted from this calculation for the 75-90° bin. To show the fluorescent intensity of Jub or Ecad at vertical or horizontal edges, for each embryo cytoplasmic fluorescence is subtracted from the average fluorescent intensity of edges in the vertical (75-90°) and the horizontal (0-15°) bins.

### Analysis of TCJs

Fluorescence within cell vertices (v) was measured using the ellipse tool in Fiji for 30-45 vertices per embryo (e). For enrichment at tricellular junctions, mean intensity of vertices was divided by the mean intensity of bicellular cell edges determined via trajectories drawn in SIESTA. For correlation analysis, ROIs of individual bicellular junctions were subjected to coloc2 in Fiji.

### Quantification of Gaps

In Fiji, on an image representing a field of view of (2136.4 µm^2^), the brush tool was used to mask each gap, and then the wand tool was used to measure area of gap and fit ellipse.

### Colocalization of signal

To determine the degree of colocalization of Cno versus α-Cat construct signal, ROIs of junctions (j) were compared for a number of embryos (e) using the coloc2 plugin on ImageJ.

### Statistics

Prism v9 (GraphPad) was used for statistical analysis and plots, except for the balloon plot in Fig. 4 which was made using Microsoft Excel. In violin plots, bold lines indicate the median, thinner lines represent interquartile ranges. For bar graphs, height of bar indicates mean, and error bars indicate standard deviation. Significance was calculated on measurements from each cell (c), junction (j) or vertex (v) for a number of embryos (e). In general, one-way ANOVA was used to determine significance between the levels of fluorescence in rescue conditions, and Two-way ANOVA was performed on measurements of planar polarity. In experiments with only two conditions, the Mann-Whitney unpaired two-tailed t-test was used. To compare terminal cuticle phenotypes, Chi-square analysis was performed on pairs of conditions. For more details per experiment please refer to Supplemental Table 1.

## Acknowledgements

We thanks Ritu Sarpal and Limin Wang for providing the data shown in Fig. S1. We thank Mark Peifer for discussion of unpublished data and comments on the manuscript. We thank Nick Brown, Thomas Lecuit, Mark Peifer, the Bloomington Drosophila Stock Center, the Drosophila RNAi Screening Center at Harvard Medical School, and the Developmental Studies Hybridoma Bank for reagents. We like to thank the Imaging Facility of the Department of Cell and Systems Biology, University of Toronto, for support. This work was funded by a project grant from the Canadian Institutes for Health Research (to U.T.). U.T. is a Canada Research Chair for Epithelial Polarity and Development.

## Author contributions

The project was conceived and experiments were designed by L.S. and U.T. L.S. carried out experiments and data analysis. U.T. provided supervision and raised funds. The paper was written by L.S. and U.T.

## Competing interest

The authors declare no competing interests.

## Data and material availability

All data are available in the manuscript or supplementary materials.

## Supplementary materials

Supplemental Figures S1 – S5

Supplemental Table S1

## Notes

### Competing Interest Statement

The authors have declared no competing interest.

