## Supplemental materials for "The function of α-Catenin mechanosensing in tissue morphogenesis"

Sep. 8, 2021

**Supplemental Figures and Table**

**Figure S1. Most follicular epithelia are normal when the M region is deleted.**

**(A)** Follicular epithelium clones positively marked with GFP in indicated genotypes. α-Cat null mutant cells show strongly reduced levels of Arm and display cytoplasmic aggregates of α-Spectrin. Expression of α-CatR and α-CatR-ΔM in α-Cat mutant cells substantially rescue these defects. Scale bar, 20 μm.

**(B)** Comparison of rescue activities of α-CatR and α-CatR-ΔM when expressed in α-Cat mutant cells in the follicular epithelium. Data are presented as mean±s.e.m.

**Figure S2. *α-Cat-RNAi* knockdown and distribution and activity of M region deletion constructs.**

**(A)** *α-Cat-RNAi* expressed by mat-GAL4 causes loss of any detectable cortical α-Cat in stage 8 embryos. Scale bar, 100 μm.

**(B)** Quantification of average levels of cortical α-Cat per embryo. Significance calculated by Mann-Whitney two-tailed t-test, **** = P <0.0001.

**(C)** Localization of α-Cat constructs expressed in an *α-Cat-RNAi* background. Gastrulating embryos are immunostained for HA to detect the HA-tagged α-Cat constructs.

**(D,E)** Close ups of the mesoderm (C) and lateral ectoderm (D) of *α-Cat-RNAi* expressing embryos, showing membrane tethers and loss of AJs. Cells lose contact apically while basal membranes remain in contact.

**(F)** Example images of the terminal cuticle phenotypes of embryos expressing *α-Cat-RNAi* and the indicated M region deletion constructs. Defects are quantified in Fig 1C.

**Figure S3. Planar polarity analysis of AJ proteins in embryos expressing M region deletion constructs in an *α-Cat-RNAi* background**.

**(A)** Plots of the fold changes of the average fluorescent intensity of edges within 15° bins versus that of the 0-15°bin (representing horizontal edges, 0°=anterior posterior axis). Each dot represents the ratio derived from one embryo. Transparent pink and cyan shapes reflect control and experimental means respectively, drawn to the height of each bar for ease of comparison. Where overlapping, grey is seen, where experimental mean is reduced compared to control pink is visible, and conversely where increased cyan is visible. Significance calculated by two-way ANOVA for conditions within the same row.

**(B)** Graphs show the mean junctional fluorescent signal per embryo of Ecad along edges in the horizontal or vertical bins in *α-Cat-RNAi α-CatX* embryos. Significance given by uncorrected Fisher's LSD.For (A) and (B), **** = P <0.0001, ***= P <0.0002, **=P<0.0021, *=P <0.0332.

**Figure S4. Cno recruitment to AJs is largely α-Cat-independent.**

(A) Examples of gastrulating *α-Cat-RNAi α-CatX* embryos heat-fixed and immunostained for α-CatR or α-CatR-ΔM1 (HA-tagged) and Cno. α-CatR-ΔM1 rescued embryos show a more irregular distribution than α-CatR.

**(B)** Quantification of the colocalization between Cno and α-Cat construct. Significance calculated by Mann-Whitney two-tailed t-test. ***= P <0.0002

**(C)** Live imaged examples of Cno::YFP signal in the amnioserosa of *α-Cat-RNAi α-CatX* embryos.

**(D)** Cross-section of the lateral ectoderm of an *α-Cat-RNAi α-CatR* embryo at stage 8 showing that Cno signal is apical to the HA-tagged α-Cat construct.

**Figure S5. Phenotypic categories of embryonic cuticle defects.**

Original images shown at left, overlays in cyan at right highlight regions of secreted cuticle. Color key at right corresponds to legend used in Fig. 8A and B.

**Table S1 Summary of genotypes and quantitative analysis for all Figures**

Genotypes used for each figure panel, number of embryos, measurements analyzed, statistical tests used and P values.

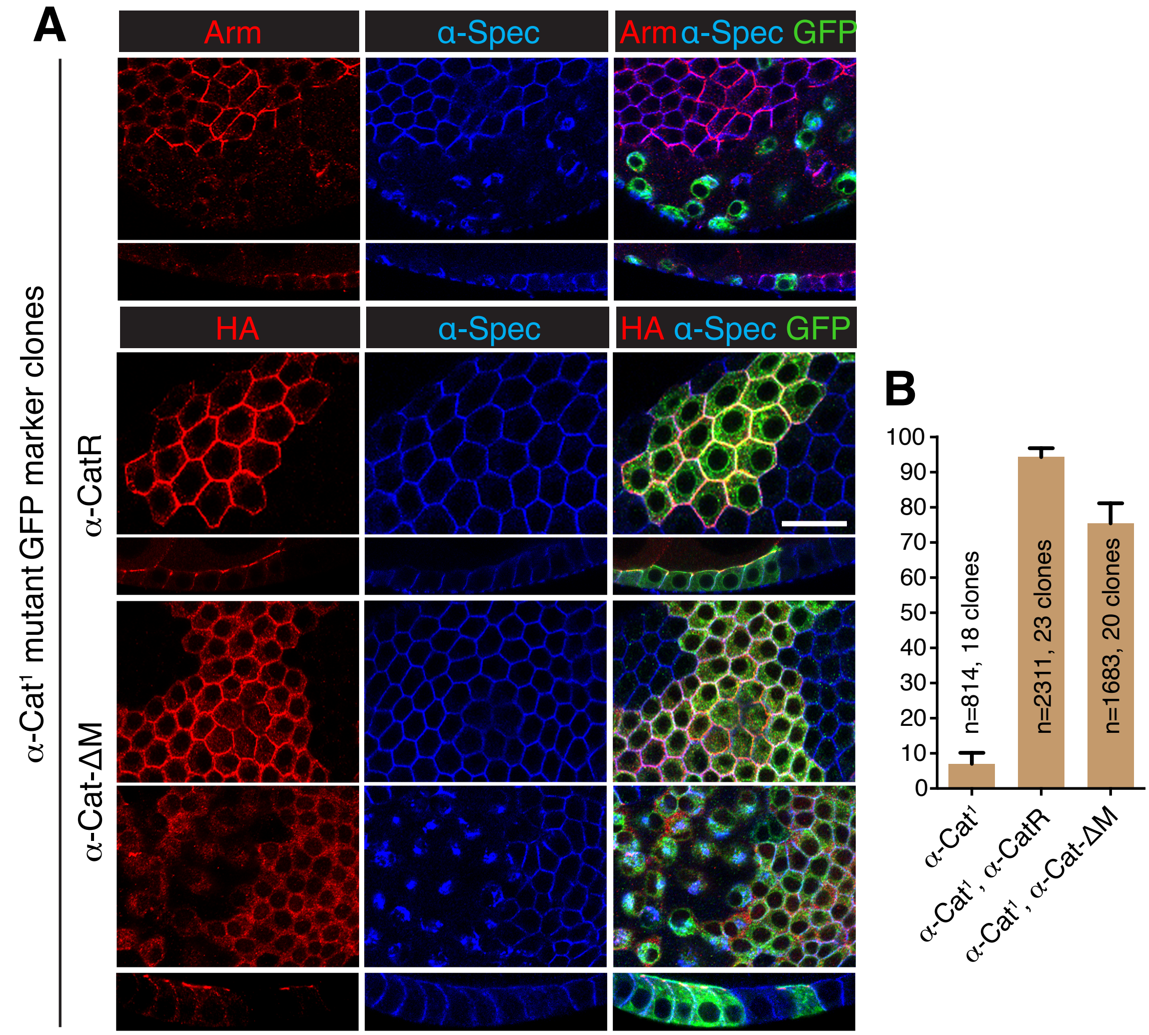


**Sheppard & Tepass, Figure S1**

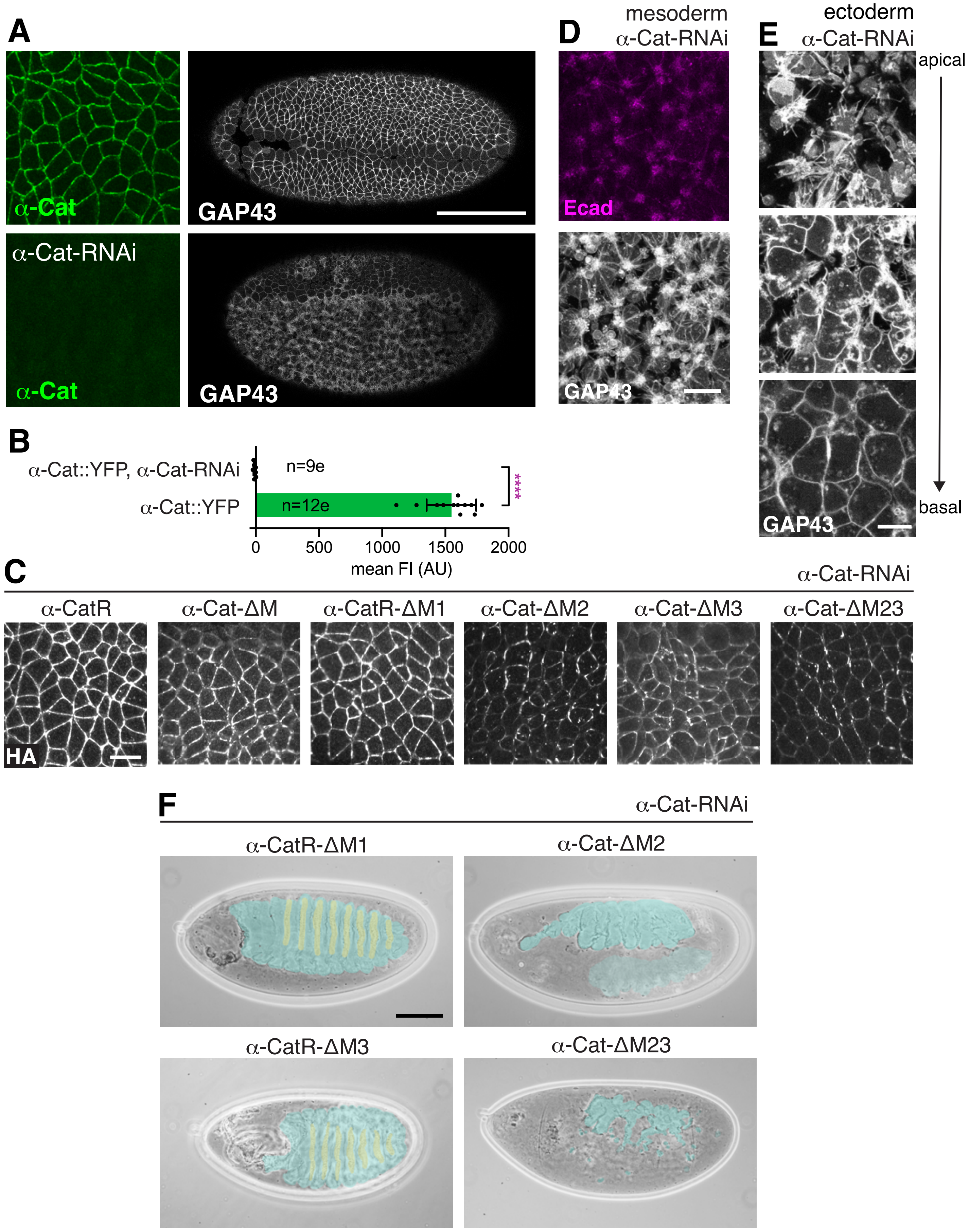


**Sheppard & Tepass, Figure S2**

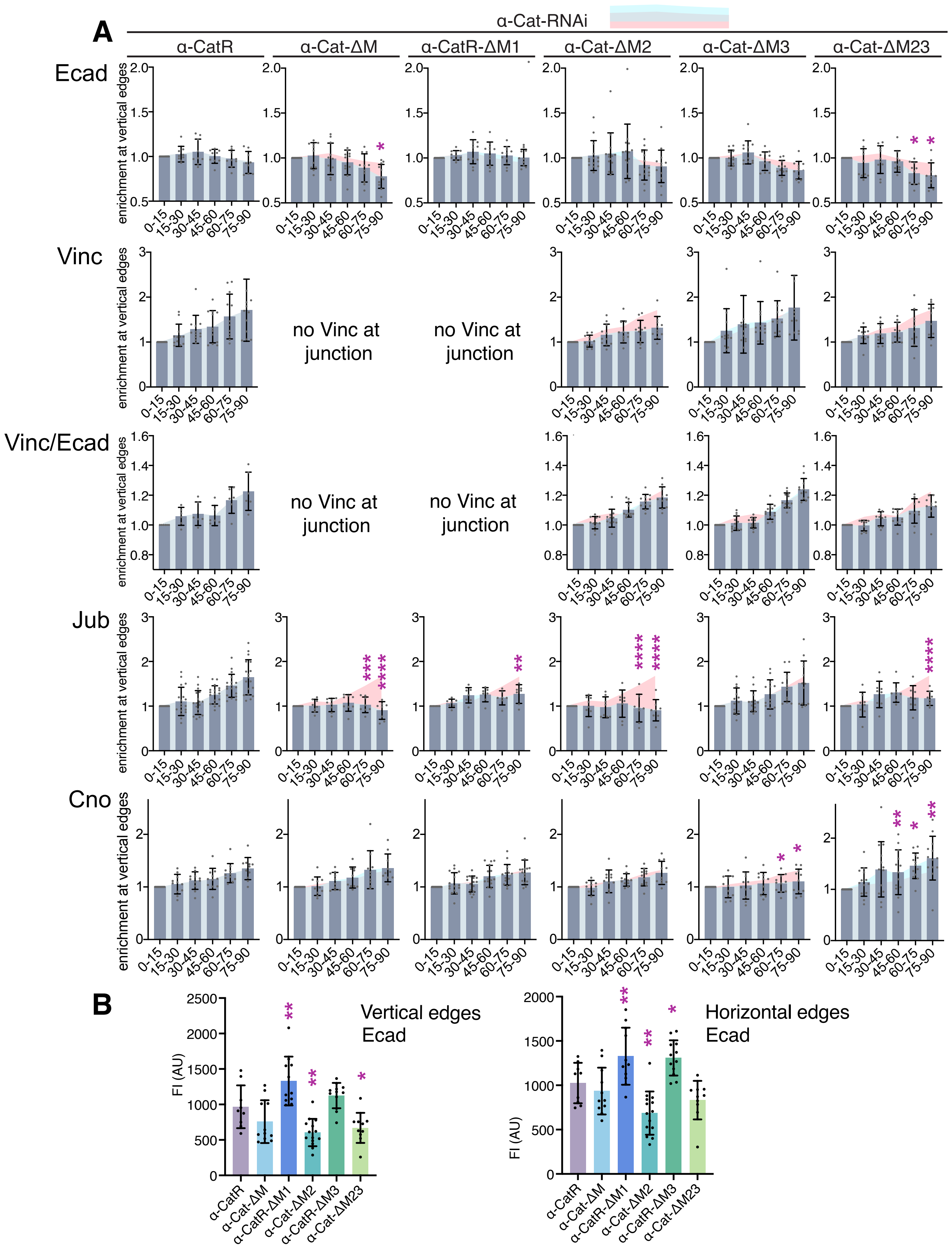


**Sheppard & Tepass, Figure S3**

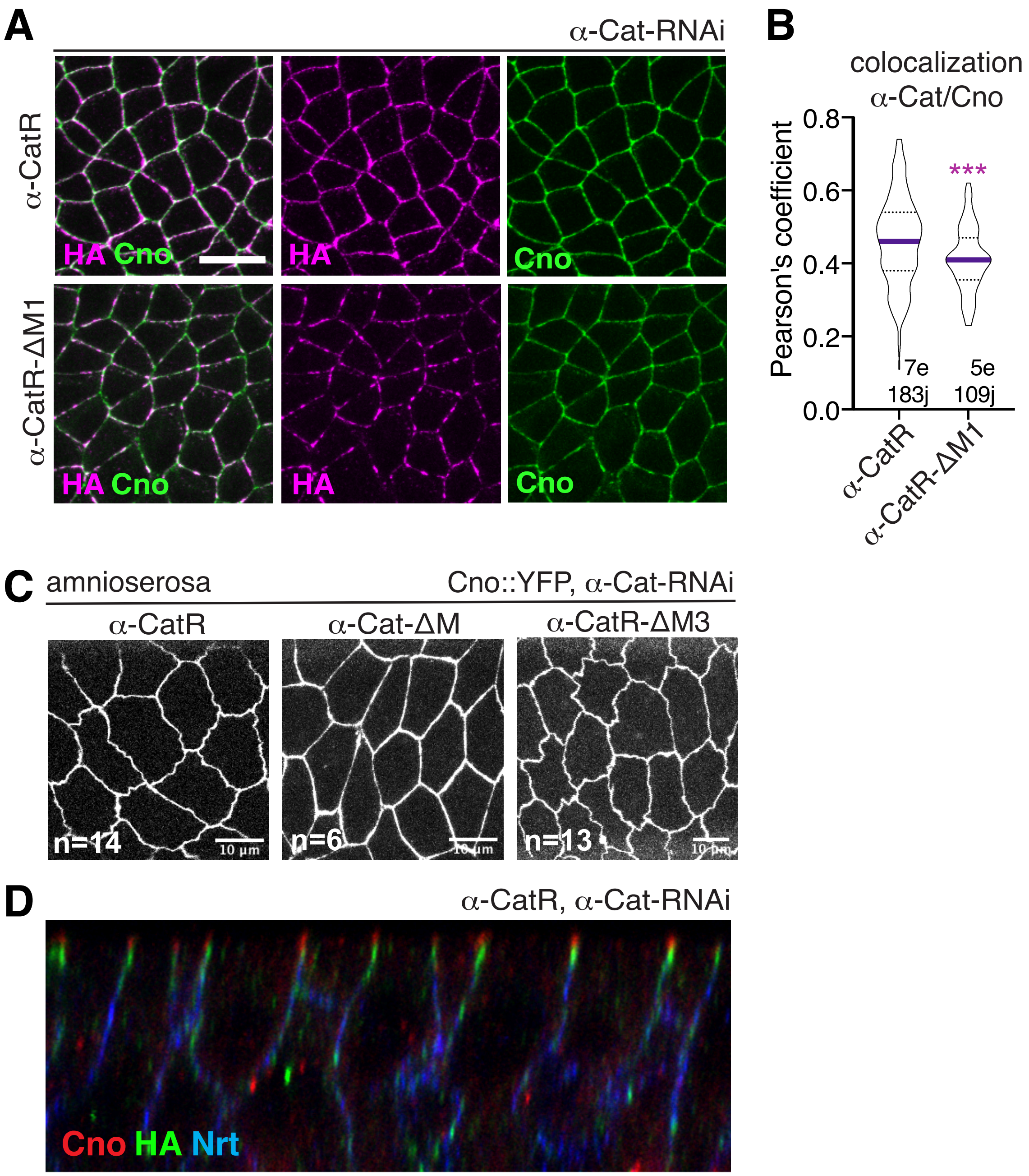


**Sheppard & Tepass, Figure S4**

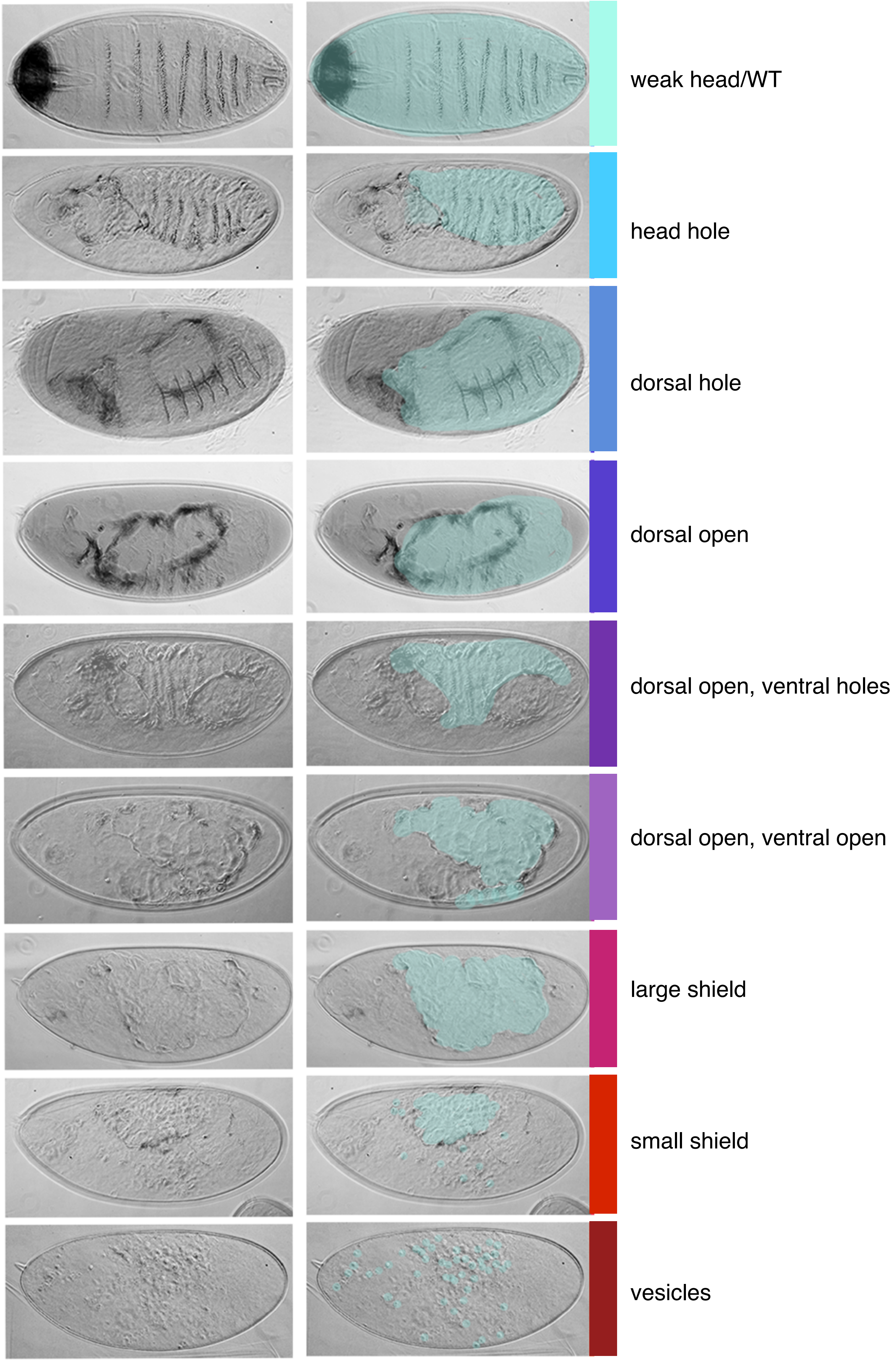


**Sheppard & Tepass, Figure S5**
